# AlphaFold2 reveals commonalities and novelties in protein structure space for 21 model organisms

**DOI:** 10.1101/2022.06.02.494367

**Authors:** Nicola Bordin, Ian Sillitoe, Vamsi Nallapareddy, Clemens Rauer, Su Datt Lam, Vaishali P. Waman, Neeladri Sen, Michael Heinzinger, Maria Littmann, Stephanie Kim, Sameer Velankar, Martin Steinegger, Burkhard Rost, Christine Orengo

## Abstract

Over the last year, there have been substantial improvements in protein structure prediction, particularly in methods like DeepMind’s AlphaFold2 (AF2) that exploit deep learning strategies. Here we report a new CATH-Assign protocol which is used to analyse the first tranche of AF2 models predicted for 21 model organisms and discuss insights these models bring on the nature of protein structure space. We analyse good quality models and those with no unusual structural characteristics, i.e., features rarely seen in experimental structures. For the ∼370,000 models that meet these criteria, we observe that 92% can be assigned to evolutionary superfamilies in CATH. The remaining domains cluster into 2,367 putative novel superfamilies. Detailed manual analysis on a subset of 618 of those which had at least one human relative revealed some extremely remote homologies and some further unusual features, but 26 could be confirmed as novel superfamilies and one of these has an alpha-beta propeller architectural arrangement never seen before. By clustering both experimental and predicted AF2 domain structures into distinct ‘global fold’ groups, we observe that the new AF2 models in CATH increase information on structural diversity by 36%. This expansion in structural diversity will help to reveal associated functional diversity not previously detected. Our novel CATH-Assign protocol scales well and will be able to harness the huge expansion (at least 100 million models) in structural data promised by DeepMind to provide more comprehensive coverage of even the most diverse superfamilies to help rationalise evolutionary changes in their functions.

## 1. Introduction

For over 30 years, the pace of sequencing proteins has outstripped that of structure determination and at the start of 2022, the non-redundant protein sequence data in UniRef90 (1) was 1000-fold greater than the associated 3D data. Methods of protein structure prediction have progressed in the same time-frame. Especially for sequences having a close homologue (>40% sequence identity) of known 3D structure, homology modelling can provide an accurate structure for many biological analyses (2–4). However, there still remains a significant deficit, including for human proteins linked to disease (5). Even for many of the well-studied prokaryotic model organisms (e.g., *Escherichia coli, Bacillus subtilis*), the proportion of the proteome (protein coding part of genome) for which high-resolution protein 3D structures have been experimentally determined or can be reliably predicted remains below 55%. The structural coverage is lower for eukaryotic model organisms, e.g., only 36% of human and 30% of baker’s yeast proteins are, at least, partially covered by structures (6).

Over the last ten years, a number of developments in template-free *ab-initio* structure prediction (e.g., co-variation (7), deep learning from vast sequence data (8)) have led to promising improvements, and recently the AlphaFold2 (AF2) method developed by DeepMind has reached an impressive level of performance as evidenced by independent assessment (9,10).

In August 2021, in collaboration with PDBe at EMBL-EBI, DeepMind provided AF2 3D-models for 21 selected model organisms (including human, mouse, *Arabidopsis thaliana*, rice, yeast and *E. coli*), comprising 365,184 model structures altogether (11). Information on global and per residue model accuracy is provided for each model. The scale and accuracy of this modelling initiative is likely to be a gamechanger for biological and medical research as protein structure data is key to understanding the molecular mechanisms of proteins and the impact of genetic variations on their molecular function and therefore the biological processes they participate in.

To exploit these data, it would be valuable to assign the modelled domains to their evolutionary families to better understand how genetic variations modify structure and ultimately function. Proteins comprise combinations of domains, and millions of combinations of domains exist across genomes (12,13). Since the protein domain is thought to be an important module contributing specific functional features, a tractable approach for handling the vast number of proteins in nature is to organise by domain family. Currently, most experimentally characterised domain structures have been assigned to fewer than 6,000 evolutionary families (14,15).

Over the last 25 years, several domain-based protein structure classifications have emerged (SCOP (16), CATH (15), SCOPe (17), SCOP2 (18), ECOD (14)) which assign experimental structures of proteins from the Protein Data Bank (PDB) to evolutionary superfamilies. ECOD and CATH are the most comprehensive, classifying 90% or more of PDB. Since the advent of large-scale genome sequencing, the CATH classification has also identified homologous domain sequences for CATH superfamilies in UniProt (19) and complete genomes from Ensembl (20). This expands the sequence population of CATH superfamilies by nearly 300-fold and brings in more functional annotations for the proteins (21).

The provision of high quality AF2 structural models from DeepMind for 21 organisms gives an opportunity to significantly expand the structural data in CATH superfamilies. This would allow us to provide multiple structural alignments and identify structurally conserved features correlating with functional motifs. It would also allow us to assess how representative existing structural superfamilies are of domains in nature and to reveal novel folding architectures and motifs not in the PDB.

Here we present a new classification protocol CATH-Assign, which incorporates novel and rapid deep learning strategies for detecting sequence and structure similarities between domains to rapidly classify structures. We applied the protocol to analyse protein structures of 21 model organisms predicted by AF2. We removed models deemed to be low quality by the AF2 developers and any that had long stretches of residues or a significant proportion of residues with no secondary structure assignment. We also removed models in which the overall contact density of residues was unusual compared to distributions seen in the PDB. Of the nearly 700,000 domains provided by AF2, only 52% of models met these criteria (369,512 domains) but 92% of them could be assigned to 3,253 existing CATH superfamilies. The remaining domains could be clustered into 2,367 putative new superfamilies. Manual curation of very remote homologues is extremely time-consuming but for a subset of these novel superfamilies, comprising human models, we manually identified 26 likely new superfamilies. By bringing the AF2 models into CATH we expand the number of ‘global fold’ groups (in which structural relatives in superfamilies superimpose well) by 36%. This suggests that the promised release of a further 100 million models by DeepMind will bring significant insights on how structures (and therefore their functions) diverge during evolution. The data will be made available grouped by CATH Superfamily and by organism through the 3D-Beacons network (22), Zenodo, and the CATH FTP.

## Results

### 2.1 Proportion of AlphaFold2 models that can be brought into CATH

#### 2.1.1 Identification of domains in AF2 protein models

We applied an in-house Hidden-Markov Model-based protocol CATH-Resolve-Hits (CRH) (23) to assign domain regions in all sequences from the 21 model organisms modelled by AlphaFold2 (AF2) (see Methods for description and Figure 2). CRH identifies domains that can be putatively assigned to either CATH superfamilies, structurally uncharacterised Pfam families or novel superfamilies (NewFams). We include structurally uncharacterised Pfam families to improve domain boundary resolution as Pfam families are manually curated and of high quality.

Using CRH and CATH (version 4.3), we assigned each region in the UniProt sequences of all AF2 models to four possible categories: CATH-experimental (CATH-PDB) if there is a structure for this domain in the PDB and predicted domains comprising CATH-HMM, Pfam or NewFam assignments. We subsequently chopped these structural regions from the corresponding AF2 protein 3D-model (See Figure 2 for the workflow and Supplementary Figure 1 for the numbers of domains in each category).

AF2 CATH-HMM models tend to be higher quality models than structurally uncharacterised Pfam or NewFam domains (see Figure 2 and Supplementary Figure 2 for more details). This is likely because AF2 provides better quality models if the structure or close homologue is present in the PDB. CATH-PDB and CATH-HMM AF2 models also have higher percentages of ordered residues (i.e., with secondary structure assignment (see Figure 1 and Supplementary Figure 2 for more details)) than the other domain types. Since AF2 model quality relies on query alignment depth, the low quality for NewFam domains could suggest significant levels of disorder or very shallow alignments for these families, with few sequences or species in them (5).

**Figure 1.**
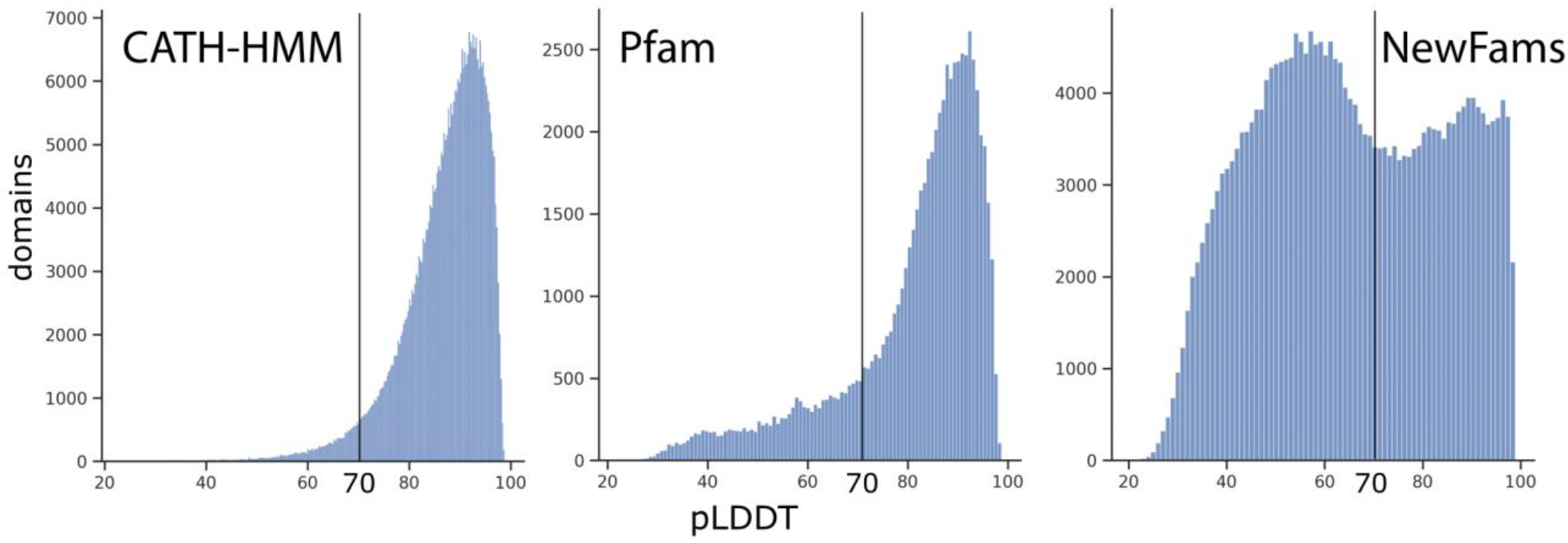
Average model quality. The plots show the distribution of average pLDDT scores for domains divided by source. The pLDDT threshold for good model quality is highlighted (>= 70).

**Figure 2.**
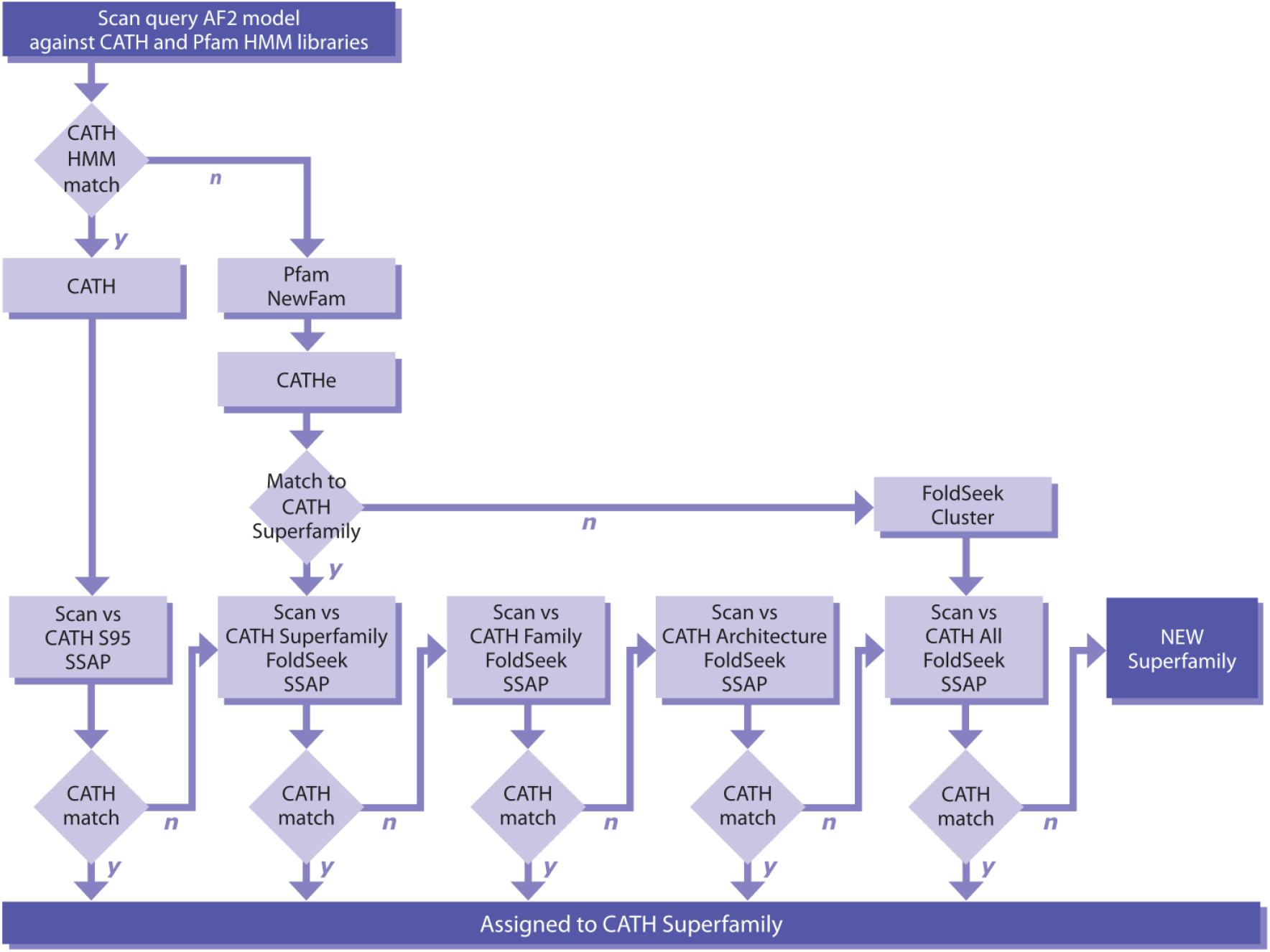
Overview of the CATH-Assign protocol. Processing of the predicted AF2 domains. CATH-HMM (labelled as CATH) are simply compared against the Superfamily non-redundant representative that they match. Pfam and NewFam domains are classified into CATH Superfamilies using the CATHe predictor where possible. A cascade method is used to validate, starting with predicted Superfamily, then architecture then everything if necessary.

#### 2.1.2 Removing models with poor quality or problematic features

Only good quality domain models (as described below) were considered for assignment to CATH superfamilies. Models that contain problematic features are likely to make it difficult to recognise a structural relationship. We used a number of criteria, including the threshold given by AF2 developers for good model quality (i.e., pLDDT >= 70) (9). We also removed models that had long regions of residues (>30% of the domain) with no secondary structure assignment or where a significant proportion of the domain (>65%) comprised residues with no secondary structure assignment (see Methods, section 4.5 to 4.8). We also imposed the criteria used for classification of domains into CATH, that domains have >= 40 residues and >= 3 secondary structure elements. Finally, we removed models in which the average contact density of the residues was very unusual compared to the values found for experimentally characterised protein domains in the PDB (see Supplementary Table 1 and 2, Supplementary Figure 3 and Methods for more details on methods used to filter the AF2 domain models and for the numbers of models removed and remaining for each type). Using these criteria, we removed 339,428 domains from further analyses.

#### Proportion of domains that can be assigned to CATH superfamilies

##### (a) Analysis of CATH-HMM predicted domains

For 273,346 domains with clear homologues in CATH (i.e., domains matching HMMs built on CATH structural representatives) we compared the domain to the structural representative used to seed the HMM. We used our slow but sensitive SSAP method to do this (see Supplementary for more details). On average, the SSAP score for the CATH-HMM domains aligned over the non-redundant (S95) representative was 84.8, with an average overlap over the S95 representative of 81.5%. A total of 246,143 CATH-HMM domains could be classified into CATH superfamilies.

##### (b) Analysis of Pfam and NewFam predicted domains

To determine whether these domains were distant homologues of CATH superfamilies, we used a recently established protein Language Model (pLM, Prot-T5 (24)) to generate protein sequence embeddings of the domains. We then used CATHe (25), an established artificial neural network (ANN) predictor trained on known CATH superfamily domain embeddings (see Methods section 4.3 for details), to predict CATH superfamilies for these uncharacterised domains. In previous studies, CATHe showed 86% accuracy on all CATH superfamilies and 98% for prediction of the 50 most highly populated superfamilies that currently account for 40% of CATH domains (25). 43.4% of Pfam domains and 23.8% of NewFam domains could be assigned to CATH superfamilies with high confidence by CATHe. We then validated the predicted domains by performing structure comparison against all non-redundant (S95) relatives in the predicted CATH superfamily.

Some superfamilies are very large (sizes of the matched superfamilies ranged from 1 to 4,546 non-redundant relatives). We therefore applied a new structure comparison method, Foldseek, which is several orders of magnitude faster than the well-established TMalign method while matching its sensitivity (26). We benchmarked this method using manually curated CATH domain assignments to determine an acceptable threshold for superfamily assignment (see Supplementary Materials, section Foldseek benchmark).

A cascading approach was used, i.e., if no match was obtained using Foldseek, our in-house SSAP method (27) was used which is very much slower but slightly more sensitive than Foldseek (See Supplementary Figure 4). Suitable SSAP thresholds were also benchmarked (see methods SSAP Benchmark). Using these approaches a total of 18,383 Pfam and NewFam domains could be classified into CATH superfamilies.

##### (c) All domains unmatched to CATH superfamilies by CATH-HMM, CATHe, Foldseek, SSAP

Finally, all unmatched domains were scanned with Foldseek and SSAP against AF2 domains assigned to CATH superfamilies by steps (a) and (b) above. This brought a further 43,646 domains into CATH superfamilies.

In summary, of the 369,512 domains passing our selection criteria and analysed using our sequence-(CATH-HMM, CATHe) and structure-based (Foldseek, SSAP) protocols (i.e., steps a-c above), 341,213 (92.3%) could be assigned to CATH superfamilies, representing a 67% expansion in CATH domains (341,213/500,238=0.67). A majority of these domains (79.3%) were relatively close homologues that could be detected by HMM based strategies (Figure 3). Table 1 shows the numbers of domains assigned to CATH superfamilies in steps (a-c) and the resulting percentage expansion in CATH.

**Figure 3.**
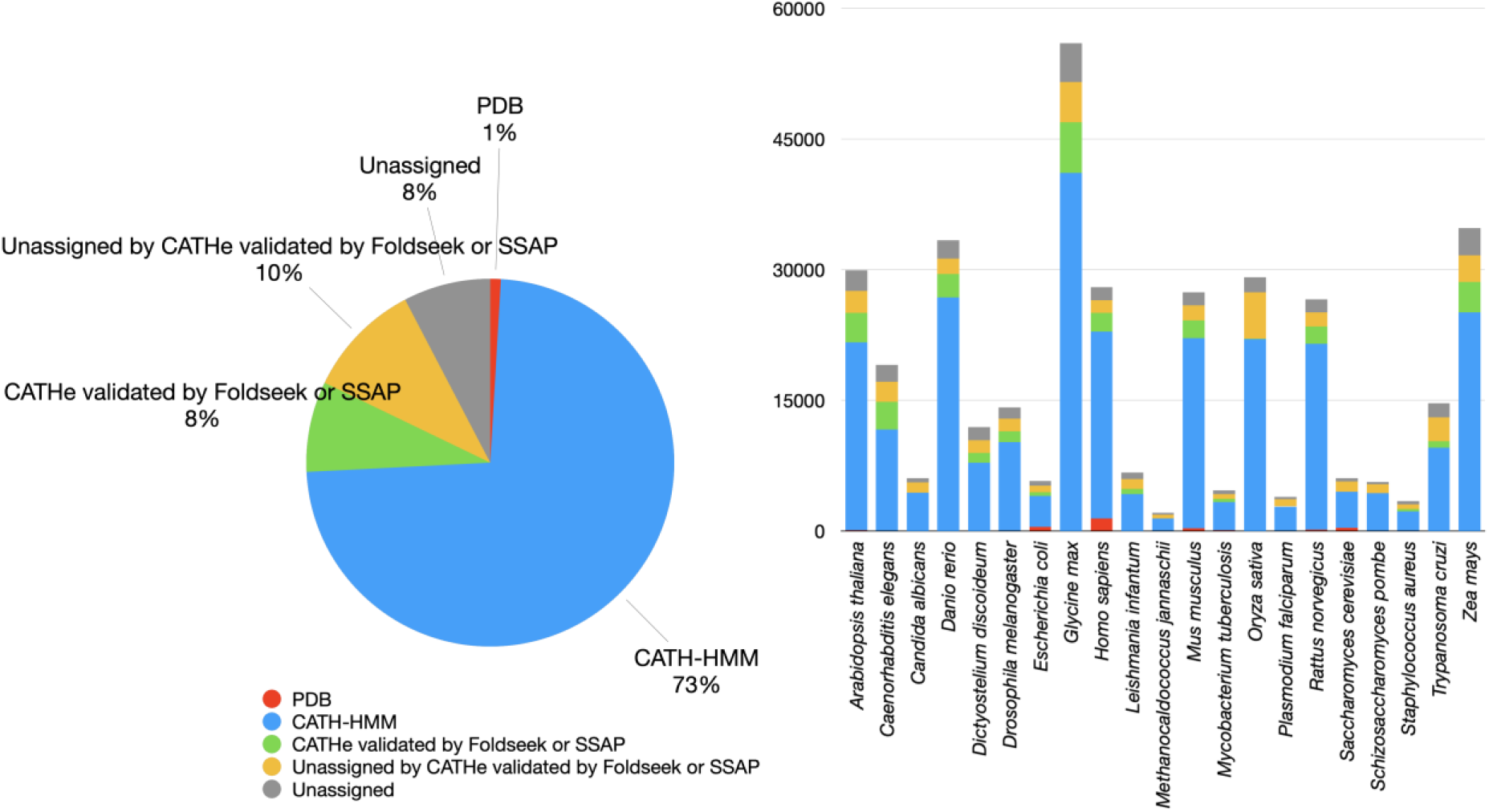
Overview of domain quality and ontology for the total AlphaFold2 dataset (left) and subdivided by each proteome (right).

**Table 1:**
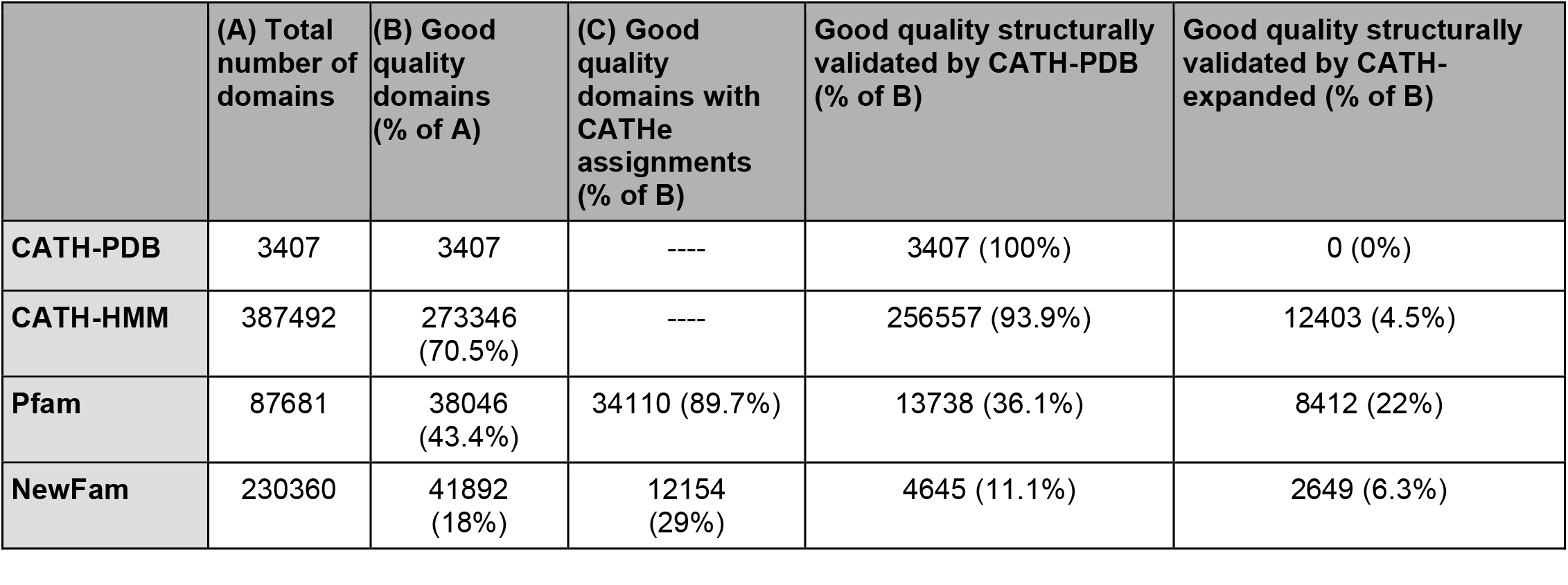
Number of domains at each processing step by source.

|  | (A) Total number of domains | (B) Good quality domains (% of A) | (C) Good quality domains with CATHe assignments (% of B) | Good quality structurally validated by CATH-PDB (% of B) | Good quality structurally validated by CATH-expanded (% of B) |
| --- | --- | --- | --- | --- | --- |
| **CATH-PDB** | 3407 | 3407 | ---- | 3407 (100%) | 0 (0%) |
| **CATH-HMM** | 387492 | 273346 (70.5%) | ---- | 256557 (93.9%) | 12403 (4.5%) |
| **Pfam** | 87681 | 38046 (43.4%) | 34110 (89.7%) | 13738 (36.1%) | 8412 (22%) |
| **NewFam** | 230360 | 41892 (18%) | 12154 (29%) | 4645 (11.1%) | 2649 (6.3%) |

These validated structures represent an average increase in structural coverage by CATH domains of 741-fold over the 21 model organisms (see Figure 4). The expanded coverage is particularly evident in the case of *Glycine max* and *Danio rerio*, with a 4,200-fold and 2,200-fold increase in structural coverage of domains belonging to these organisms.

**Figure 4.**
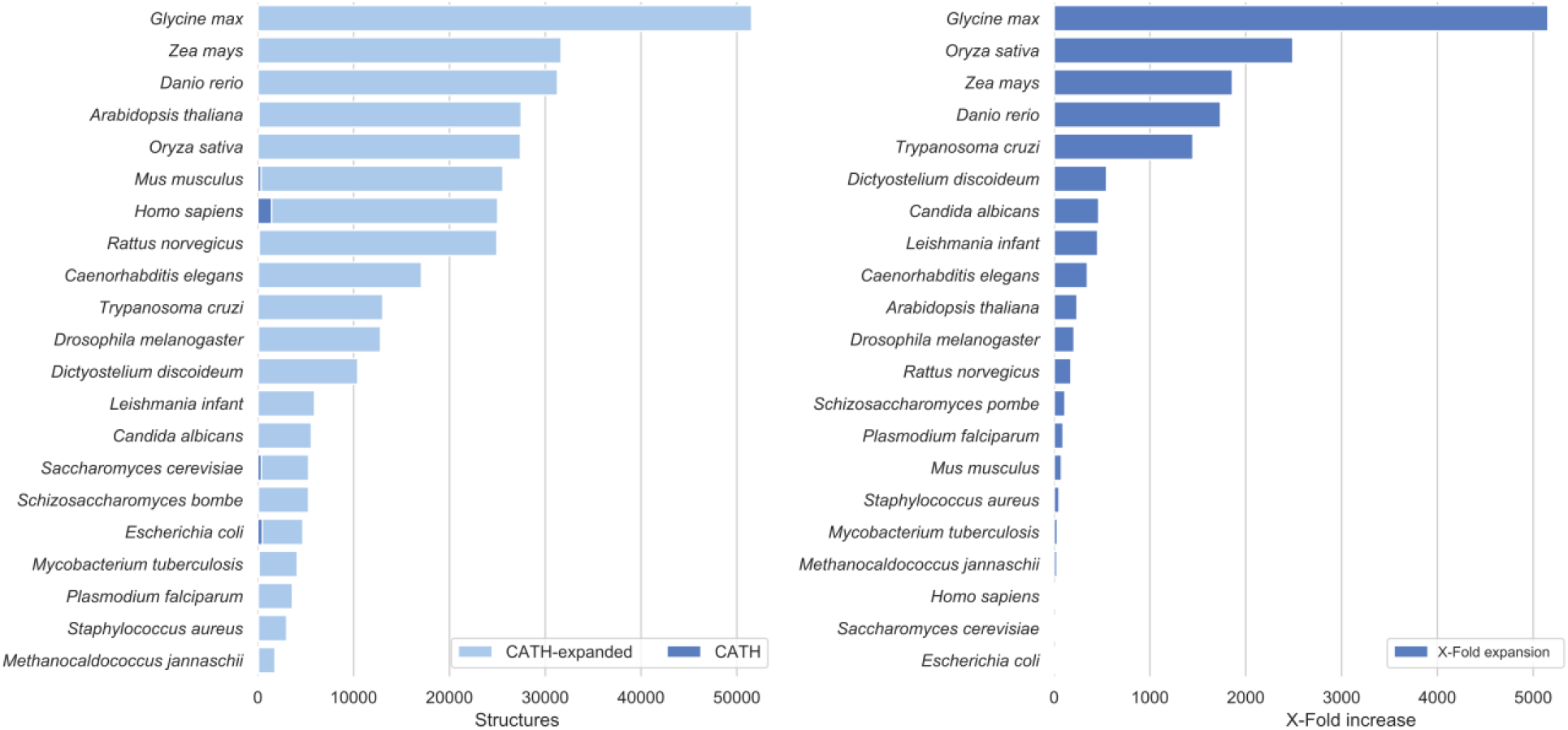
Expansion in structural coverage by total number of structural domains (left) and fold-wise (right) by validated CATH-HMM, Pfam and NewFams domains models for the 21 organisms in the AlphaFold2 dataset. The left panel showcases the total number of CATH domains derived from a protein entry from each organism (CATH), as well as CATH assignments now covered by an AF2 structure (CATH-expanded). The right panel showcases the X-fold expansion in the number of structural domains by organism.

Considering the class of the domains, 29.8% map to mainly-alpha superfamilies, 23.4% to mainly-beta and 46.7% to alpha-beta, and these proportions are quite similar to those observed for experimental domain structures in CATH, albeit with a slightly lower percentage of alpha superfamilies in CATH experimental (alpha: 21.2%, beta: 23.4%, alpha/beta: 46.7%). This overabundance of mainly-alpha superfamilies has been noticed also in other AF2 domain classification efforts based on SCOPe (17). Supplementary Figure 5 shows the average expansion of each CATH architecture. The top 200 most highly populated superfamilies in CATH (sometimes referred to as MegaFamilies) comprise nearly 70% of the experimental structures in CATH. A significant portion of AF2 domains (62%) map to these superfamilies.

### 2.5 Expansion of functional families by AlphaFold2 structural data

Each CATH superfamily has been classified into functional families (FunFams), in which relatives are predicted to have similar structures and functions (21,28,29). Since this sub-classification is computationally expensive (29), FunFams have only been generated for families in which at least one relative has an experimental characterisation in the Gene Ontology. Version 4.3 of CATH contains 212,872 FunFams comprised of 322,202 domains (i.e., 64% of all CATH domains). Only 17,208 FunFams (8%) have at least one domain which has been structurally characterised. Assignment of AF2 3D-models into CATH superfamilies by the protocol described above (Figure 2) brought 104,306 more structural relatives into the FunFams. This increased the proportion of FunFams that have at least one relative with an experimental or AF2 structure by 3.7-fold overall and by up to 6.6-fold, depending on the organism (see Figure 5). Supplementary Figure 6 shows that FunFams with higher numbers of relatives and species within them were more likely to have good quality AF2 models predicted.

**Figure 5.**
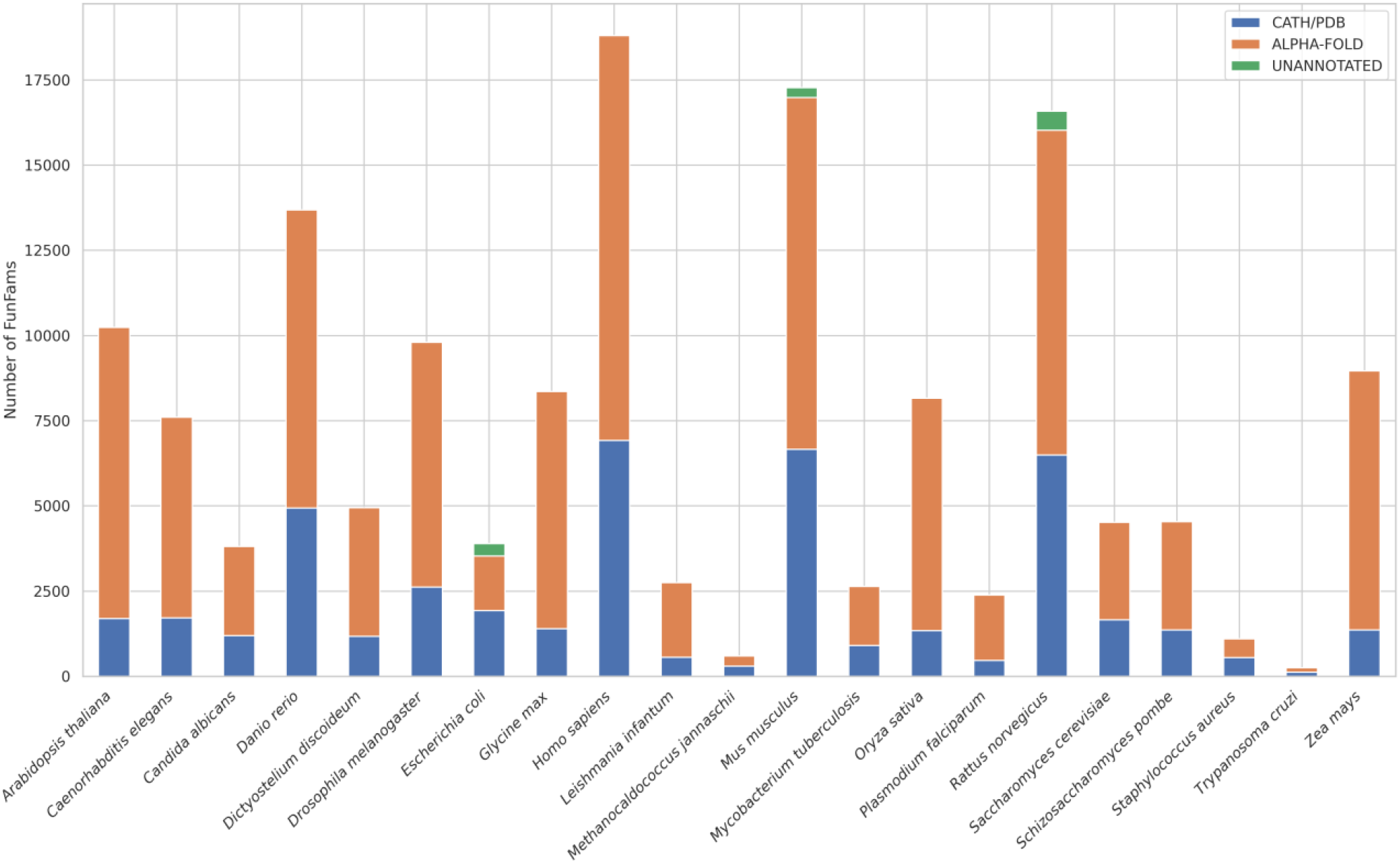
Structural coverage expansion of CATH FunFams by AlphaFold domains.

### 2.6 Identification of new superfamilies and new folds

To assess whether the remaining 34,502 unclassified AF2 domains belonged to novel superfamilies, we compared the domains against each other using Foldseek and then SSAP, and structurally similar domains were clustered together (see Methods for more details), giving 7,519 structural clusters. For those clusters where relatives had been assigned to a superfamily by CATHe a representative was structurally compared against CATH-expanded domains (i.e., PDB domains and classified AF2 domains) in the CATH fold group predicted by CATHe. If no match was found it was scanned against the CATH architecture predicted by CATHe. This was done as manual analyses of selected CATHe results showed good performance in predicting fold and architecture. If no confident CATHe assignment was obtained, we scanned against all domain structures in CATH-expanded. Scans were first performed by Foldseek and if no match found then SSAP was used (see Figure 2 for more details).

Using this approach we assigned a further 6,203 domains to CATH superfamilies, from 2,899 clusters. Of the remaining 4,620 clusters for which there was no structural match to CATH superfamilies, 1,154 matched structures in the PDB not yet classified in CATH. A further 714 were clearly multidomain as we found using Foldseek that part of the region structurally matched a domain in CATH-expanded. For the remaining 2,367 detailed manual analysis is required to check for very remote homology to the closest matched CATH superfamily relatives (30). This is time consuming as it requires extensive visual inspection and checking information in the literature. Furthermore, since we had observed that bringing AF2 domain models into CATH improved the detection of remote homologues by providing greater coverage of the family (see section 2.1.2(c) above), we reasoned that detection of very remote homologues would be better enabled once further AF2 models (more than 100 million for UniRef90 promised in 2022) had been processed and brought into CATH.

We therefore opted to manually examine the 618 putative new families/folds containing at least one relative from *Homo sapiens*, to assess the nature of new families and folds and assess whether there were any interesting and unusual structural features or perhaps previously missed problematic features that could explain why they had not been matched to existing CATH superfamilies/folds. We examined a representative from each of the 618 structural clusters.

For 89% of the clusters, we found possible explanations for the lack of structural matches with domains in CATH superfamilies. Some of these cases (42%) appear to be regions comprising more than one domain (see Figure 6C). This was most frequent for domains assigned to structurally uncharacterised Pfam families, for which the lack of structure can make it difficult to determine domain boundaries. A further 23% appeared to contain problematic regions e.g., large unstructured regions at either termini or poorly packed secondary structures (see Figure 6A, B and D) not picked up by the thresholds on our filtering programs, but which would make it hard to recognise structural similarities with relatives in CATH. Some of these features may reflect inaccurate models caused by the small family sizes available to AF2 to model the structures.

**Figure 6.**
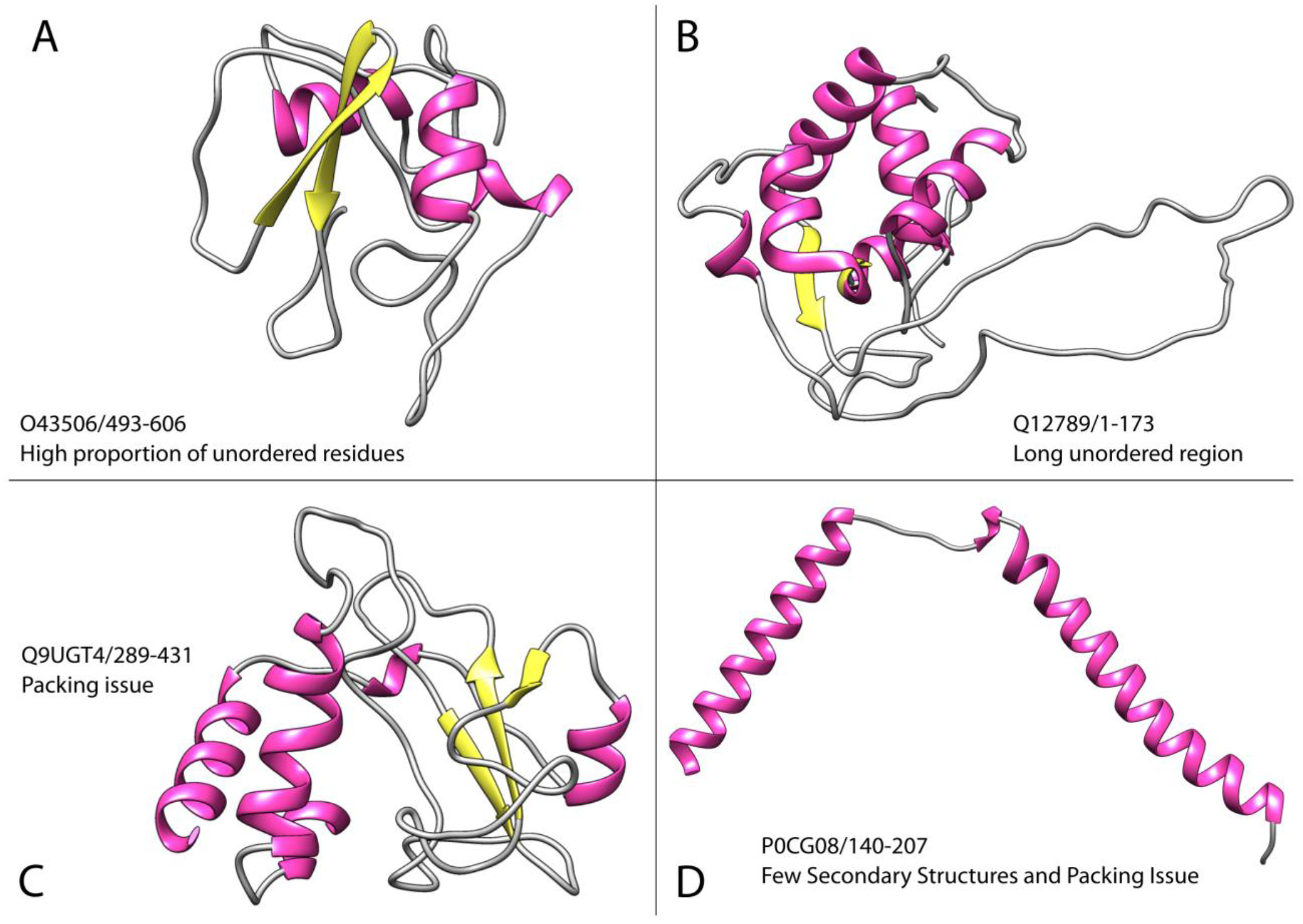
Issues encountered when processing domains not assigned to CATH. Each structure figure was generated using UCSF Chimera (31), with identifiers in the format UniProt_ID/start-stop.

There were 26 putative clusters remaining representing putative new superfamilies (see Supplementary Figures 7 and 8). Some looked similar to known families in CATH and it is likely that with the release of additional AF2 models for UniRef90 and subsequent expansion in the coverage of CATH with these models, these putative novel domains will match a CATH AF2 relative. Some have unusual structural architectures. For example, to our knowledge there are no examples of alpha-beta propellors (see Figure 7C) in any of the domain structure classifications (i.e., SCOP, CATH, ECOD). Even more unusual is the heart shaped arrangement (Figure 7D) that may comprise three small repeat domains (small alpha beta 2-layer domains) linked by longer helices. This protein, whose existence was confirmed also by RosettaFold (32), is a mitochondrial T-cell activation inhibitor involved in T-cell activation and memory formation and we found other related AF2 domain structures in 14 species. We were also able to build a dimer with AlphaFold Multimer (33) and in complex with two interactors (CLIC3, MAN1A1) predicted by STRING (34). This is not a compact globular domain as there is no clear hydrophobic core. However, it is hard to see how individual domains could be carved out of this. It is an unusual multidomain protein as in most multidomain proteins the domains are typically closely packed. It will make more sense once the functional role and structures of the protein partners are known.

**Figure 7.**
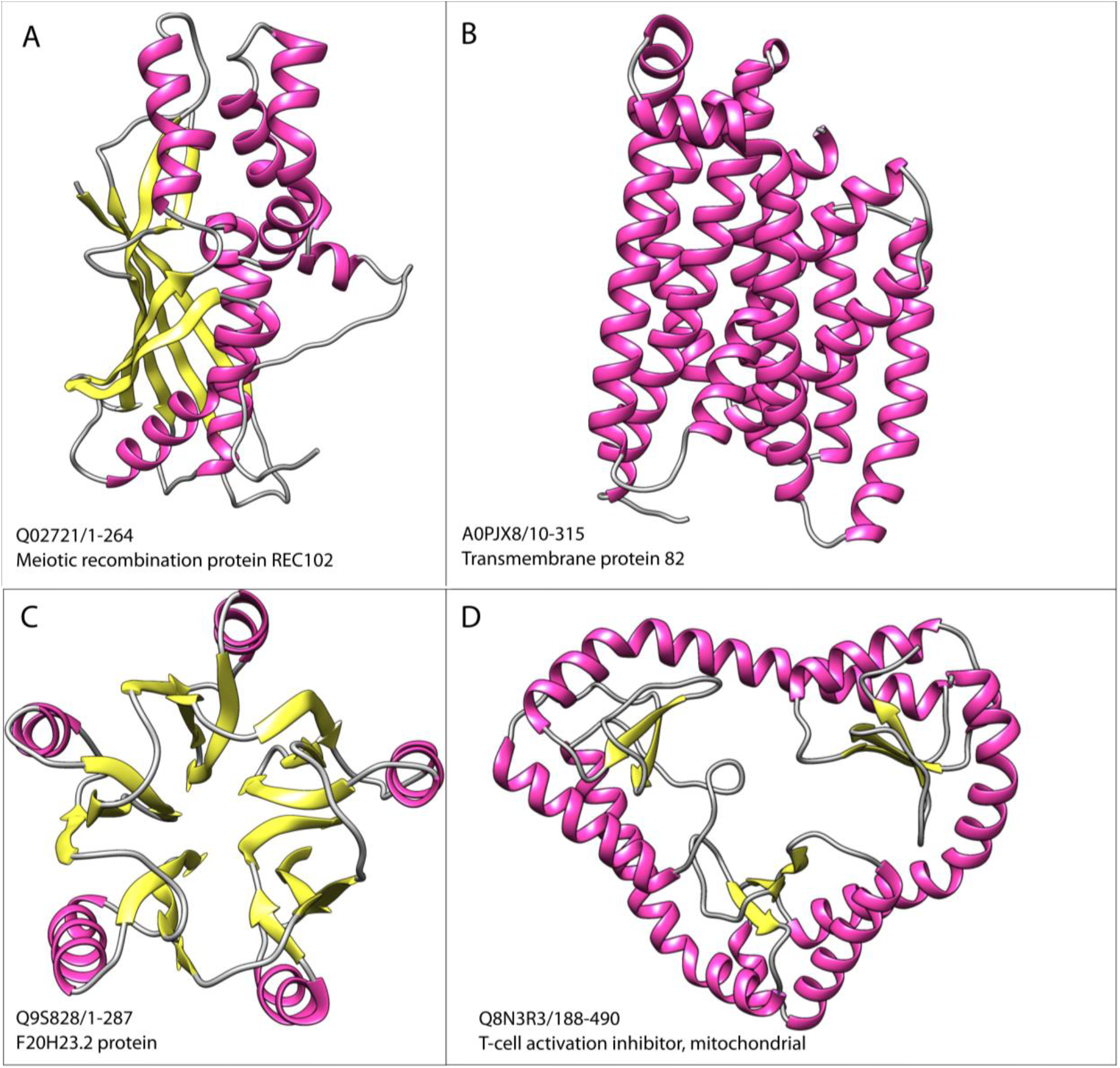
New Structural Superfamilies. Each structure figure was generated using UCSF Chimera, with identifiers in the format UniProt_ID/start-stop.

We will revisit these putative novel human superfamilies and those from the other model organisms once DeepMind releases the UniRef90 models to see if expanding CATH further helps to bring some of them into existing CATH superfamilies as extremely remote homologues.

Although ∼92% of the good quality AF2 models can be assigned to existing structural superfamilies in CATH, they bring considerable structural novelty. CATH superfamilies are sub-clustered into groups of relatives that can be superposed well (structurally similar groups (SSGs)). These can be considered as ‘global fold’ groups as there are distinct changes outside the common structural ‘core fold’ (see Figure 8). Currently there are ∼28,000 such global fold groups in CATH. Adding AF2 structures increases this number by ∼36% to 38,000.

**Figure 8.**
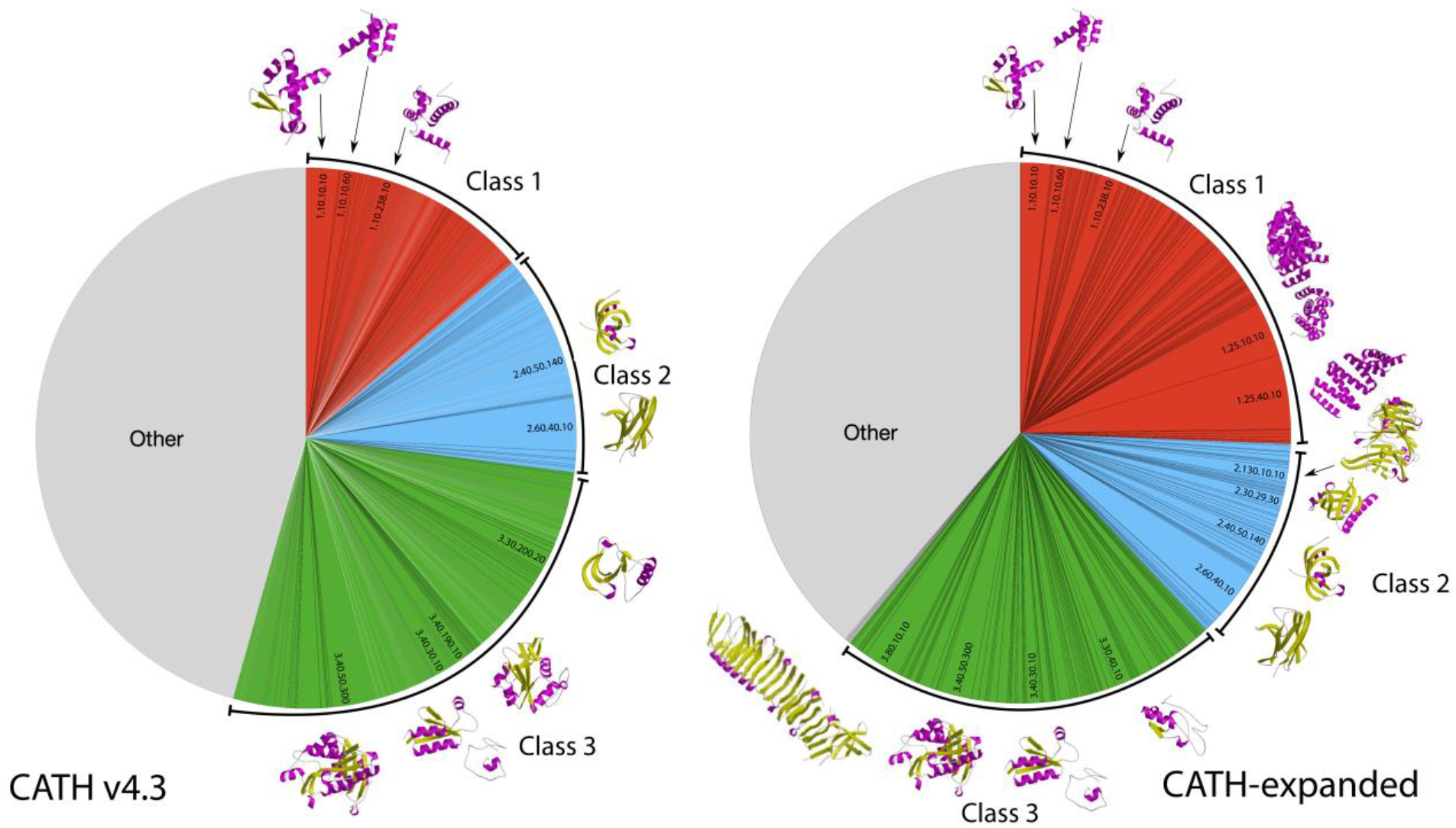
Structural diversity in CATH Superfamilies (left) and expanded by AF2 models (right).

## 2. Discussion

For the last 50 years, the number of structures known experimentally has been a small proportion of the numbers of protein sequences determined. Just prior to the release of DeepMind’s AlphaFold2 data, this discrepancy was more than 1000-fold if metagenome sequences are also considered. The validation of AF2 by CASP14 (35) has given confidence in the quality of the AF2 structural models, and the promise of nearly 100 million AF2 models (for UniRef90) by the end of 2022 represents a landmark in the study of protein structure and protein evolution.

One of the biggest immediate challenges is to handle the scale of the data. The promised expansion in the data represents a 200-fold increase in the number of protein structures. Traditionally, structure comparison algorithms have been much slower than sequence comparison methods. However, the launch of AF2 occurred almost in parallel with the development and release of an extremely fast new comparison method, Foldseek (26). Foldseek has comparable accuracy to the TM-align method (36), traditionally used to assess structural similarity by many biologists and also employed by the CASP evaluation committee, but is 20,000 times faster. In addition, related machine learning tools to those employed by AF2, such as our CATHe predictor, that exploit language models to capture information about the structural contexts of residues, have become widespread and are being applied in the field of protein family classification as they are more powerful and faster than HMM based approaches (24,25,37–39). These new technologies will harness the information from AF2 to enhance our understanding of fold space and the structural mechanisms by which structural changes impact on the functions of proteins.

In this work we have built on classification workflows developed over the last 25 years for classifying protein domains in the CATH database <u>(20)</u>. These include strategies for identifying domain regions in protein sequences (CATH-HMM, CATH-Resolve-Hits (23)) and strategies for detecting very remote homologies by sensitive structure comparison methods (SSAP (27)). To handle the scale of the AF2 models we built the CATH-Assign protocol, which used these approaches together with an extremely fast structure comparison method (Foldseek) and also a novel protein language model (ProtT5 (24) developed by the Rost group) in a classifier (CATHe (25)) for detection of extremely remote homologues (<20% sequence identity) (see Figure 2).

We were able to process 369,512 good quality AF2 models in less than six months. Since CATH-Assign is now established, new releases of AF2 models will be processed much faster. The near 70% expansion in structural data in CATH is impressive for such a short time scale. Prior to AF2, CATH contained nearly half a million experimental structures, in 5,660 superfamilies. Classification of these superfamilies was performed over 25 years and benefited from substantial manual curation. Since coverage of CATH superfamilies by experimental structures is extremely sparse (on average <5%) structural validation can be difficult for very remote homologues. However, the pending release of 100 million AF2 models for UniRef90, whilst requiring significant effort to process, will significantly facilitate validation and curation as structural coverage of superfamilies will increase as we bring more AF2 models in. Another problem caused by the sparse coverage of CATH superfamilies is that domain boundary detection by CRH is likely to be less accurate for AF2 domains very distant from anything in CATH, as sizes of CATH relatives could very different from the domain in the AF2 protein. This increases the difficulty of assigning these AF2 domains to CATH superfamilies, as very stringent thresholds on domain overlap are used to verify homologues. This is particularly problematic for structurally uncharacterised Pfam domains assigned to AF2 models by CRH, as the Pfam boundaries will not have been verified structurally and any errors will be transferred by CRH. However, as imported AF2 models gradually expand structural coverage of CATH superfamilies, most AF2 domains will match homologues in CATH, and we will be able to replace CRH with Foldseek. We are also exploring in-house and publicly available deep learning predictors for *ab-initio* domain boundary assignment.

We remove nearly 50% of predicted domain models from our analyses, depending on the organism (see Supplementary Figure 9). This is in agreement with early studies that calculated the percentage of disordered and low-quality regions in AF2 models in *Homo sapiens* (40). The percentage of discarded domains is higher for Eukaryotes, with the exception of yeasts (baker’s and fission) (Supplementary Figure 10). A large proportion (52%) of these have poor model quality (pLDDT < 70) and residues not predicted as ordered (i.e., not in secondary structures according to DSSP (41)) (3%). We also removed domains containing long continuous regions with no secondary structure, LURs, (>= 30% of the residues in the protein), as it would be difficult to match these domains to the more ordered protein domains in CATH (only 2.7% of CATH domains have LURs). It is not clear whether these regions are disordered or regions where AF2 struggles to model the conformation of the residues. They may also represent regions that undergo conformational change on binding to other proteins and therefore capable of adopting multiple confirmations. We found quite a large number (79,825, 23%) of predicted domains with less than three secondary structures. From manual inspection, many comprise well-predicted alpha helices which were often not packed against each other or domains in the proteins (see Figure 6D for an example). It would be interesting to seek sequence relatives across diverse species for evidence of conserved residues suggesting functional roles.

Our results give an interesting perspective on the structural models in the 21 model organisms. CATH-Assign brought 92.3% of AF2 good quality models (having no problematic features) into CATH superfamilies. Our study manually evaluated putative new superfamilies in human and identified 26 novel superfamilies. Although our results suggest another 3,081 new superfamilies in the other 20 organisms, as discussed many may match AF2 models pulled into CATH in the future. It is also likely that AF2 relatives pulled into CATH will include ‘bridging’ relatives that allow us to merge CATH superfamilies.

Taking into account, the proportion of removed problematic models, CATH could expand by 50-fold or more once the new UniRef90 models are brought in. Development of CATH-Assign and the establishment of stringent and well benchmarked thresholds for HMM, CATHe, Foldseek and SSAP matches puts CATH in a good position to process this data in a timely way.

We will continue to refine our characterisation of the novel superfamilies and exploit the expanded structural coverage of CATH superfamilies to further probe the relationships between structure and function. CATH-Assign will be applied to further releases of AF2 models. The CATH AF2 domains will be made available grouped by CATH Superfamily and by organism through the 3D-Beacons network (22), Zenodo, and the CATH FTP.

In summary, recent developments in deep learning methods applied to the analysis of protein structures (i.e., Foldseek) and protein sequence (i.e., pLMs exploited for classification - CATHe) have enabled the rapid processing of 708,941 predicted domain models generated by AF2. Although nearly 48% of domains from the model organisms were removed (because they were poorly modelled or had features that made them problematic for structure comparisons to globular domains in CATH), of the remaining domains 92% could be assigned to one of 5,600 CATH superfamilies. We identified 3,081 putative novel superfamilies. We manually examined a subset of 618 of these found in human and identified 26 which currently appear to be novel. The small number of new superfamilies identified to date is perhaps not surprising. In contrast, the expansion in structural diversity in CATH superfamilies (i.e., 36% increase in global fold groups) brought by AF2 relatives is exciting, as it could help rationalise functional divergence.

## 3. Methods

### 4.1 3D Models retrieval and processing

A total of 365,184 3D-models for 21 model organisms modelled with AlphaFold2 (AF2) were retrieved from the AlphaFold Protein Structure Database FTP (https://www.alphafold.ebi.ac.uk/) (9,11). Due to a tool crashing on non-ATOM records, all models were stripped of all non-ATOM records. The sequences of the AF2 models are based on reference proteomes from UniProt, therefore MD5 hashes for each protein sequence in the reference proteomes were generated to facilitate tracking of pre-existing annotations, mapping of unique domain sequences and to avoid differences in naming across CATH, UniProt reference proteomes and AlphaFold DB structures.

### 4.2 MD5 hashes and domains previously assigned by CATH-HMM and Pfam

For each unique MD5s in the dataset we assigned predicted protein domain boundaries. Gene3D assigns CATH domain annotations to UniProt entries by scanning their sequences against a library of 62,915 Hidden Markov Models seeded by a structural representative from each cluster of CATH relatives (at 95% sequence identity)(42).

Existing annotations and their boundaries from CATH and Pfam were retrieved from Gene3D and used as input for CATH-Resolve-Hits (CRH)(23). CATH-Resolve-Hits assigns the best possible combination of domains for a protein sequence to obtain the optimal coverage. A region in each protein could be therefore assigned to a CATH domain, a Pfam domain or to a domain-sized region unassigned to either of those (dubbed ‘NewFams’) (Figure 2). In our protocol we used a 40-residues criterion to recognise domain sized regions as this threshold has been used to populate CATH.

### 4.3 CATH superfamily predictions for Pfam and NewFams domains

We assigned a tentative, to-be-validated, superfamily CATH code for each domain sequence (Pfam and NewFams) using CATHe, a deep-learning based method for detecting remote homologues for CATH superfamilies. The first step in the CATHe pipeline was to convert the sequence domains from CATH-HMM matches into a numerical representation (sequence embedding) using the ProtT5 protein Language Model (pLM). The pLM provides residue level embeddings which are then mean-pooled to obtain the embedding for the entire protein sequence. An Artificial Neural Network model was employed to learn from these sequence embeddings and to predict the superfamily annotations for new sequence domains. CATHe was trained on 1773 CATH superfamilies and attained a prediction accuracy of 85.6% +-0.4% (95% confidence interval) on them. In order to use this model to make new predictions, we conducted a threshold analysis from which we concluded that a 40% prediction probability (correlating to an error rate of 5%) was optimal for our use case. Domain assignments below the 5% error rate threshold were marked as unassigned.

### 4.4 Domain chopping from AlphaFold2 models

Predicted domains were chopped from the AF2 models using a built-for-purpose Python pipeline based on the pdb-selres module from pdb-tools (43). The algorithm uses CATH-Resolve-Hits (CRH) output files or CATHe predictions and performs some initial checks, such as a lookup of the MD5 for the corresponding proteome and assigning the MD5 to a UniProt entry.

AF2 uses multiple models for large proteins over 2700 residues, providing 1400 amino acids long, overlapping fragments that are shifted by 200 residues. Based on the predicted domain boundaries, the algorithm detects which fragment contains the full domain, chops and creates a new PDB file with additional headers with metadata such as the domain MD5, the file from which it was chopped and if available, the assigned Pfam family or predicted CATH superfamily.

### 4.5 Chopped domain quality assessment

Each domain quality was assessed by calculating the average pLDDT score. The quality of the domain was calculated as the average pLDDT of the constituent residues of the domain.

### 4.6 Long Unordered Regions

Using pLDDT-per-residue scores, we identified long unordered regions (LUR) as regions at least five residues long with a pLDDT < 70. Domains with more than 30% of residues in a single LUR were discarded.

### 4.7 Secondary Structure Elements and Order Predictions

The secondary structure of each domain was assessed using DSSP (41), with the resulting files optimised on secondary structure element lengths by secmake (https://github.com/UCLOrengoGroup/secmake). The DSSP predictions were used for secondary structure elements assignments and filtering. The overall unordered prediction was calculated as the percentage of residues not part of secondary structure elements over the total number residues. Domains with more than 65% of residues unordered were removed from our classification analysis.

### 4.8 Packing density and Globularity Predictions

We used two metrics to predict the globularity of the domains obtained from the AF2 structures. The first one predicts the packing density by calculating the average number of neighbour residues each hydrophobic residue in the protein has within 5 Å. This was done using the python Bio.PDB package (44). While this first metric considers the chemical aspect of it, the second metric predicts globularity on a mechanical level, by calculating the surface and volume of the domain. Using the program MSMS included in PyMOL (45), we obtained both the solvent excluded surface (SES) area as well as the volume resulting from it. For the metric we then calculate the quotient of SES area / Volume. The more globular a protein is, the smaller should this value be.

To obtain thresholds for both metrics we ran them on the set of domain structures within CATH, which have been hand-curated over the years. There are a total of 61,238 domains in this dataset, which we constrained to only alpha, beta and mixed alpha-beta proteins. In order to account for errors in the dataset, we took as a globularity threshold the top 95% of hits in the dataset. This results in a packing density of 9.75, and a SES area / Volume value of 0.494 (Supplementary Figure 3).

### 4.9 Superfamily assignment validation protocol

We created a library comprising non-redundant representatives (at 95% sequence identity - S95s) for all 6331 superfamilies in CATH.. We performed an initial scan of all query domains against the library of S95 representatives for the predicted superfamily using Foldseek, developed by the Steinegger Group (https://github.com/steineggerlab/foldseek), with a minimum overlap set at 0.4, coverage mode based on query and sensitivity set at 9. After extensive benchmarking (see Supplementary section Foldseek Benchmark), we set an overlap threshold of 60% and bitscore thresholds of 106 (for classes 1 and 3) and 165 (for class 2) for superfamily recognition. All queries with hits above the threshold were set aside, while the remainder were run against the same library using the in-house SSAP using established thresholds for homology (Supplementary section SSAP Benchmark) (27).

### 4.10 Clustering of unassigned domains into tentative new Superfamilies

All domains for which no superfamily could be assigned were scanned in an all-vs-all fashion using Foldseek with an overlap set at 60%, bitscore of 165 (the strictest for any given CATH class, see Supplementary section Foldseek Benchmarking) and sensitivity set at 9. The resulting output file was then fed into TCluster (46,47) ran with single linkage clustering using the bitscores as weights for clusters cut-off.

## Acknowledgments

NS, VN, IS, CR and VW acknowledge funding from the BBSRC grants BB/R009597/1, BB/S020144/1, BB/V014722/1, BB/R014892/1, BB/T002735/1 and BB/S020039/1. NB acknowledges funding from the Wellcome Trust grant 221327/Z/20/Z. M.S. acknowledges support from the National Research Foundation of Korea grants [2019R1-A6A1-A10073437, 2020M3-A9G7-103933, 2021-R1C1-C102065, 2021-M3A9-I4021220]; and the Creative-Pioneering Researchers Program through Seoul National University. S.K. acknowledges support by the National Research Foundation of Korea (NRF) grant No. 2019R1A6A1A10073437. SV acknowledges funding support from the European Molecular Biology Laboratory – European Bioinformatics Institute. This work was additionally supported by the Bavarian Ministry of Education through funding to the TUM and by a grant from the Alexander von Humboldt foundation through the German Ministry for Research and Education (BMBF: Bundesministerium für Bildung und Forschung), by two grants from BMBF (031L0168117 and program “Software Campus 2.0 (TUM) 2.0” 01IS17049) as well as by a grant from Deutsche Forschungsgemeinschaft (DFG-GZ: RO1320/4-1).

We thank the DeepMind Team for making the data (and associated methods) available to the research community and the EBI Teams for establishing the AlphaFoldDB portal.

## Notes

### Competing Interest Statement

The authors have declared no competing interest.

