## Supplementary Materials for "AlphaFold2 reveals commonalities and novelties in protein structure space for 21 model organisms"

### Supplementary Materials and Methods

### Contents

**Supplementary Table 1.** Number of domains at each step of processing.

**Supplementary Table 2.** Number of domains discarded by reason.

**Supplementary Figure 1**: Predicted domains in AlphaFold DB by source.

**Supplementary Figure 2.** Distribution of model quality (top) and percentage of residues not in secondary structures (bottom) by domain source and CATH class.

**Supplementary Figure 3.** Scatter plot of packing density and Surface Area / Volume with marginal distributions for the protein domains in the CATH database. The dashed lines show the 95% cutoff for each metric, which has been used to label the AlphaFold domains as globular or non-globular.

**Supplementary Figure 4.** Total proportion (left) of domains structurally validated using CATH-PDB and CATH-expanded by Foldseek and SSAP, and by organism (right).

**Supplementary Figure 5**. CATH Architecture expansion by AF2 models.

**Supplementary Figure 6**. AF2 domains model qualities in FunFams versus sequences in each FunFam.

**Supplementary Figure 7.** All alpha/beta novel folds in AF2.

**Supplementary Figure 8.** All mainly-alpha novel folds in AF2.

**Supplementary Figure 9.** Distribution of AF2 domains not included in CATH (left) by organism (right).

**Supplementary Figure 10.** Distribution of AF2 domains with good model quality and discarded (left) by organism (right).

**Benchmarking SSAP and Foldseek for homologs detection**

**Supplementary Figure 11:** Foldseek bitscore plotted against the structural alignment overlap. Each pair of comparisons was coloured according to their homology.

**Supplementary Figure 12:** Error rate by Foldseek bitscore for each CATH class. The horizontal blue line represents the 5% error threshold.

**Supplementary Figure 13:** SSAP score plotted against the structural alignment overlap calculated as 100% x overlap /length of largest domain. Each pair of comparisons was coloured according to their homology.

**Supplementary Figure 14:** Error rate by SSAP score for each CATH class. The horizontal blue line represents the 5% error threshold.

| Species | Chopped | Filtered | Structurally validated | Percentage of domains brought into CATH over total |
|---|---|---|---|---|
| **Arabidopsis thaliana** | **54586** | **29959** | **27603** | **92.1%** |
| **Caenorhabditis elegans** | **34581** | **19057** | **17134** | **89,9%** |
| **Candida albicans** | **8509** | **6069** | **5611** | **92.5%** |
| **Danio rerio** | **66965** | **33398** | **31306** | **93.7%** |
| **Dictyostelium discoideum** | **23647** | **11916** | **10455** | **87.7%** |
| **Drosophila melanogaster** | **27928** | **14187** | **12883** | **90.8%** |
| **Escherichia coli** | **7315** | **5727** | **5190** | **90.6%** |
| **Glycine max** | **107848** | **56035** | **51556** | **92%** |
| **Homo sapiens** | **59314** | **28029** | **26484** | **94.5%** |
| **Leishmania infantum** | **13520** | **6700** | **5940** | **88.7%** |
| **Methanocaldococcus jannaschii** | **2513** | **2090** | **1857** | **88.9%** |
| **Mus musculus** | **55270** | **27403** | **25915** | **94.6%** |
| **Mycobacterium tuberculosis** | **6515** | **4685** | **4247** | **90.7%** |
| **Oryza sativa** | **56618** | **29116** | **27431** | **94.2%** |
| **Plasmodium falciparum** | **7187** | **3934** | **3654** | **92.9%** |
| **Rattus norvegicus** | **52663** | **26620** | **25105** | **94.3%** |
| **Saccharomyces cerevisiae** | **8526** | **6085** | **5683** | **93.4%** |
| **Schizosaccharomyces pombe** | **7618** | **5627** | **5350** | **95.1%** |
| **Staphylococcus aureus** | **4409** | **3442** | **3078** | **89.4%** |
| **Trypanosoma cruzi** | **28392** | **14650** | **13056** | **89.1%** |
| **Zea mays** | **75017** | **34783** | **31675** | **91.1%** |

**Supplementary Table 1.** Number of domains at each step of processing.

| Species | pLDDT < 70 | Domain residues not in secondary structure > 65% | Packing issues | LUR > 30% | SSE < 3 |
|---|---|---|---|---|---|
| **Arabidopsis thaliana** | **12767** | **1391** | 3858 | **981** | **5630** |
| **Caenorhabditis elegans** | **8369** | **509** | 2557 | **568** | **3521** |
| **Candida albicans** | **1109** | **91** | 523 | **107** | **610** |
| **Danio rerio** | **12976** | **1496** | 6869 | **893** | **11333** |
| **Dictyostelium discoideum** | **7083** | **313** | 1396 | **404** | **2535** |
| **Drosophila melanogaster** | **6810** | **515** | 2842 | **475** | **3099** |
| **Escherichia coli** | **243** | **43** | 464 | **63** | **775** |
| **Glycine max** | **28300** | **1685** | 8323 | **2041** | **11464** |
| **Homo sapiens** | **13590** | **1243** | 6891 | **683** | **8878** |
| **Leishmania infantum** | **4425** | **150** | 1155 | **187** | **903** |
| **Methanocaldococcus jannaschii** | **84** | **12** | 147 | **12** | **168** |
| **Mus musculus** | **11641** | **1170** | 6242 | **648** | **8166** |
| **Mycobacterium tuberculosis** | **482** | **55** | 522 | **83** | **688** |
| **Oryza sativa** | **19157** | **617** | 3397 | **729** | **3602** |
| **Plasmodium falciparum** | **2139** | **61** | 549 | **116** | **388** |
| **Rattus norvegicus** | **11245** | **1096** | 5435 | **673** | **7594** |
| **Saccharomyces cerevisiae** | **1134** | **75** | 516 | **106** | **610** |
| **Schizosaccharomyces pombe** | **779** | **70** | 486 | **94** | **562** |
| **Staphylococcus aureus** | **241** | **10** | 213 | **29** | **474** |
| **Trypanosoma cruzi** | **9519** | **341** | 2089 | **369** | **1424** |
| **Zea mays** | **24371** | **935** | 6180 | **1375** | **7373** |
| **AlphaFold** | **176464** | **11878** | 60654 | **10636** | **79796** |

**Supplementary Table 2.** Number of domains discarded by reason.

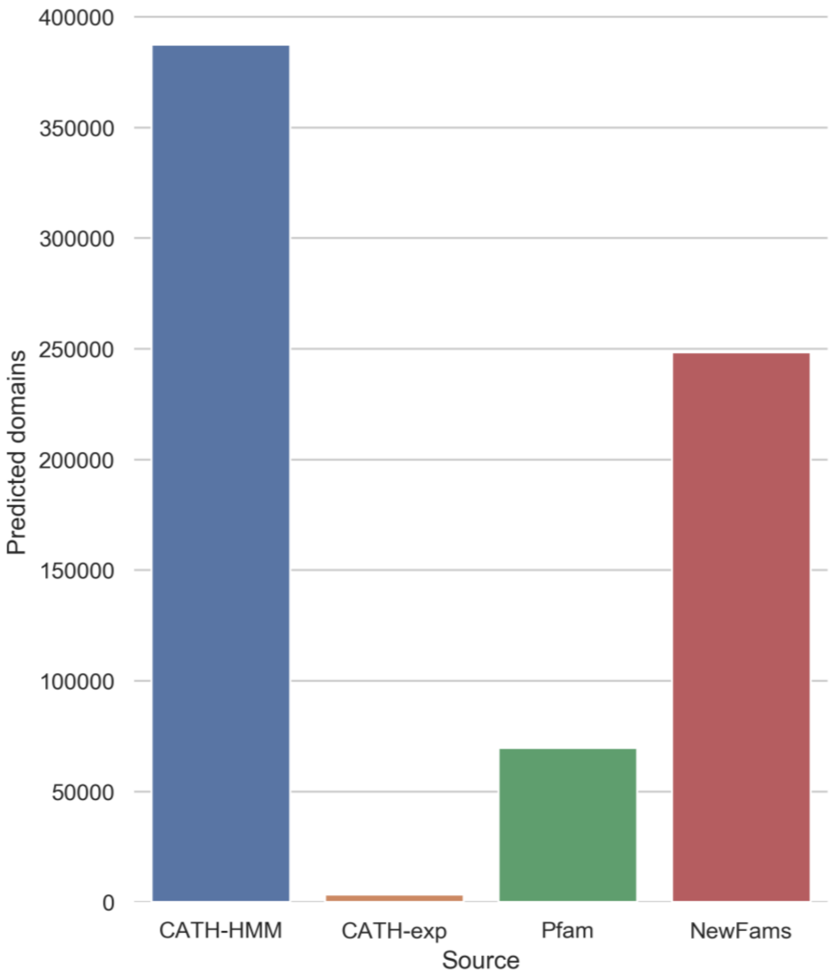


**Supplementary Figure 1**: Predicted domains in AlphaFold DB by source.

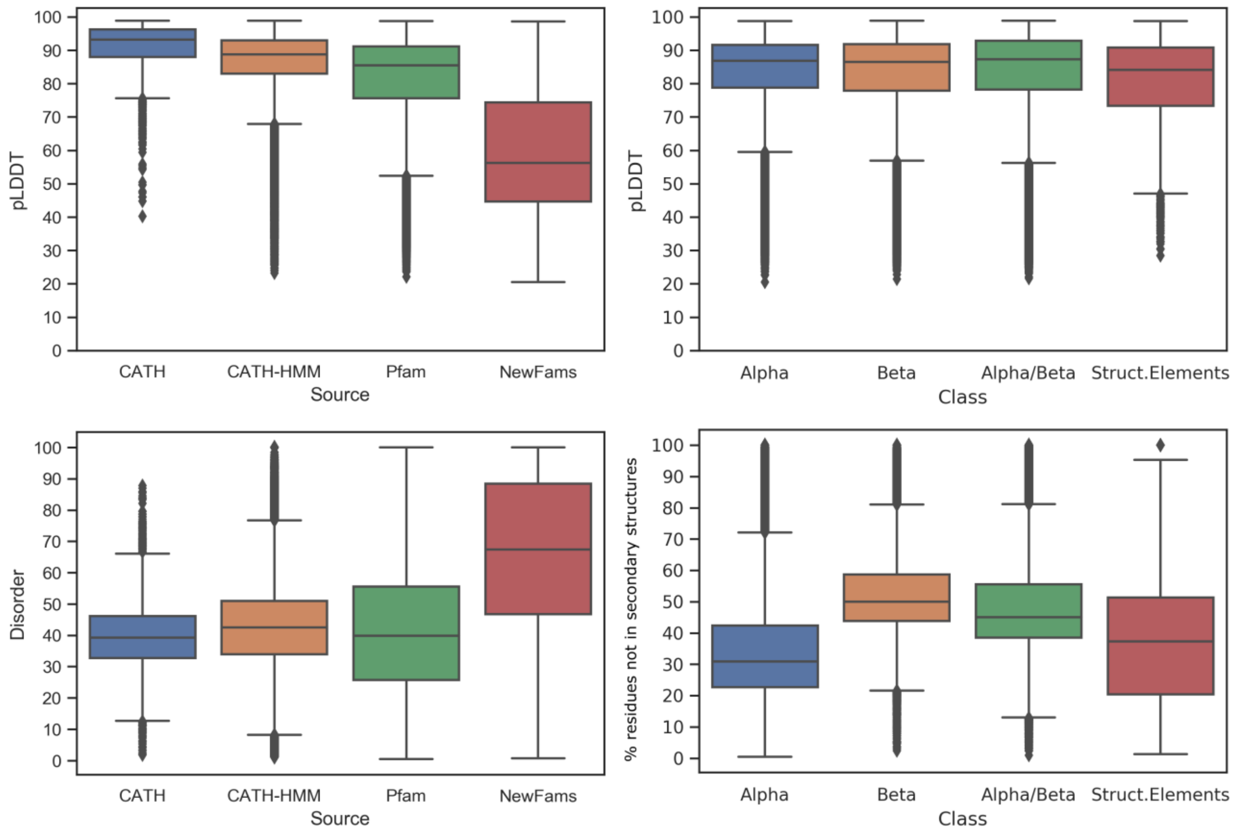


**Supplementary Figure 2.** Distribution of model quality (top) and percentage of residues not in secondary structures (bottom) by domain source and CATH class.

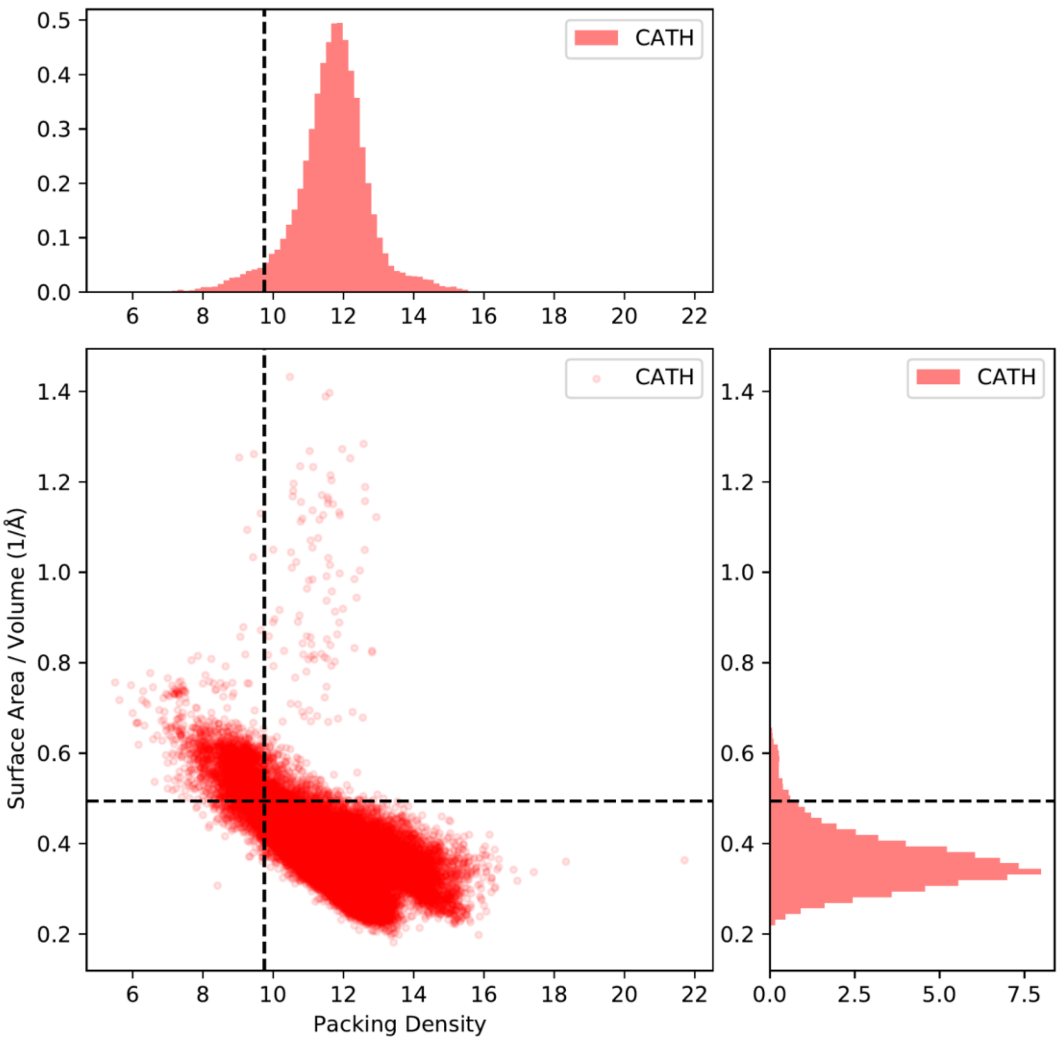


**Supplementary Figure 3.** Scatter plot of packing density and Surface Area / Volume with marginal distributions for the protein domains in the CATH database. The dashed lines show the 95% cutoff for each metric, which has been used to label the AlphaFold domains as globular or non-globular.

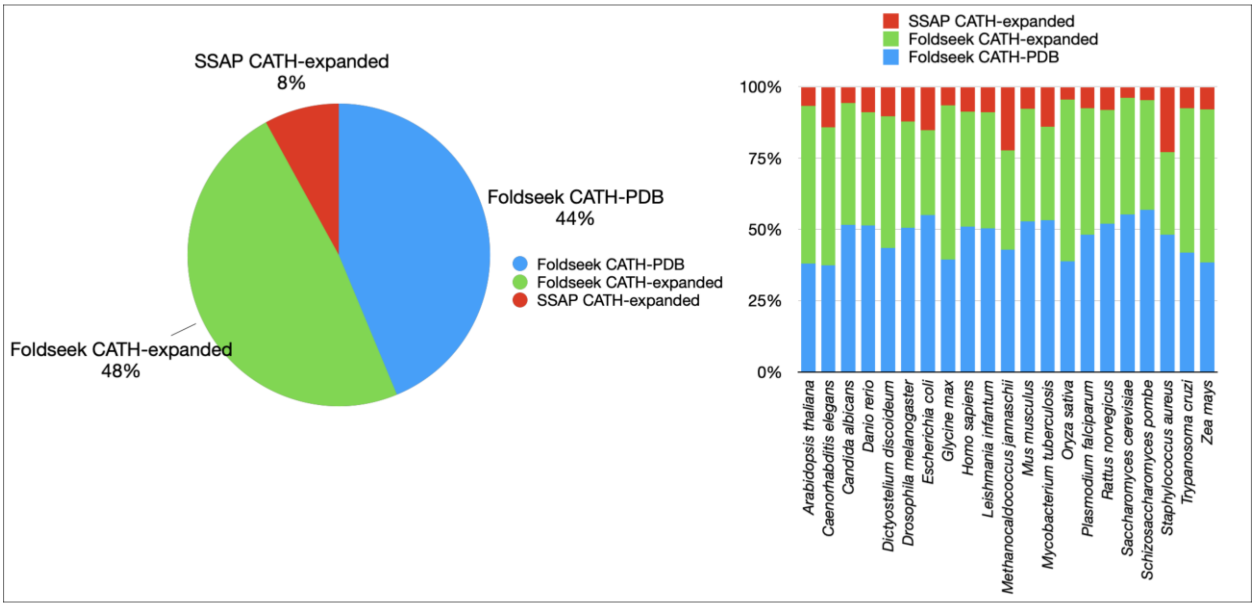


**Supplementary Figure 4.** Total proportion (left) of domains structurally validated using CATH-PDB and CATH-expanded by Foldseek and SSAP, and by organism (right).

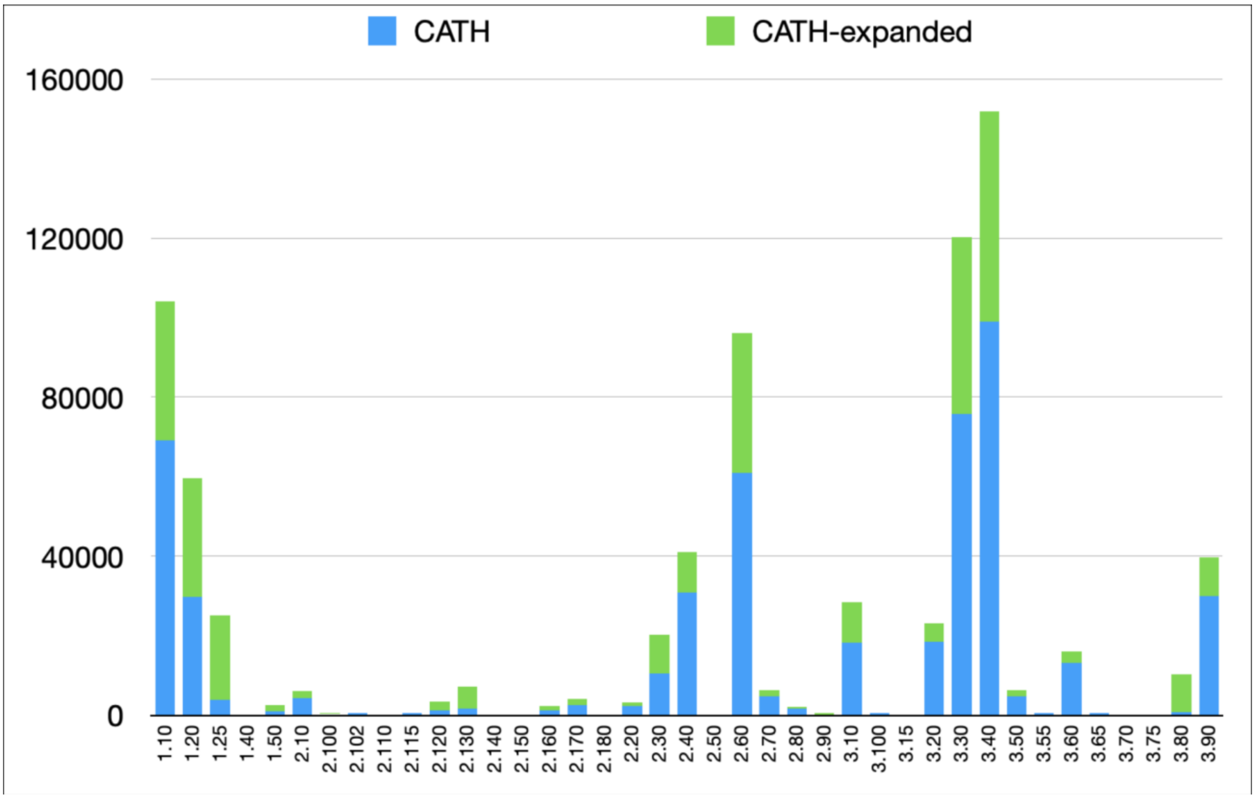


**Supplementary Figure 5**. CATH Architecture expansion by AF2 models.

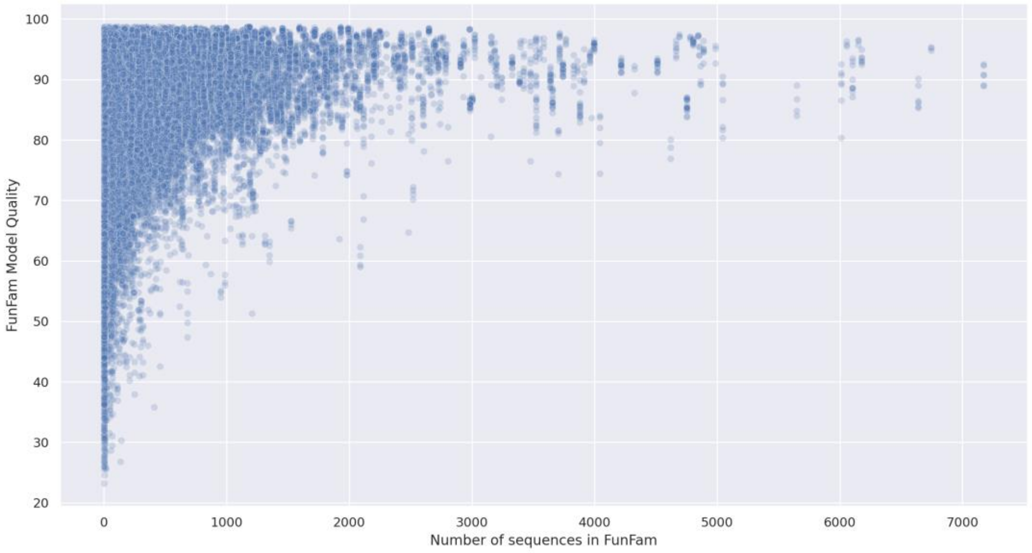


**Supplementary Figure 6**. AF2 domains model qualities in FunFams versus sequences in each FunFam.

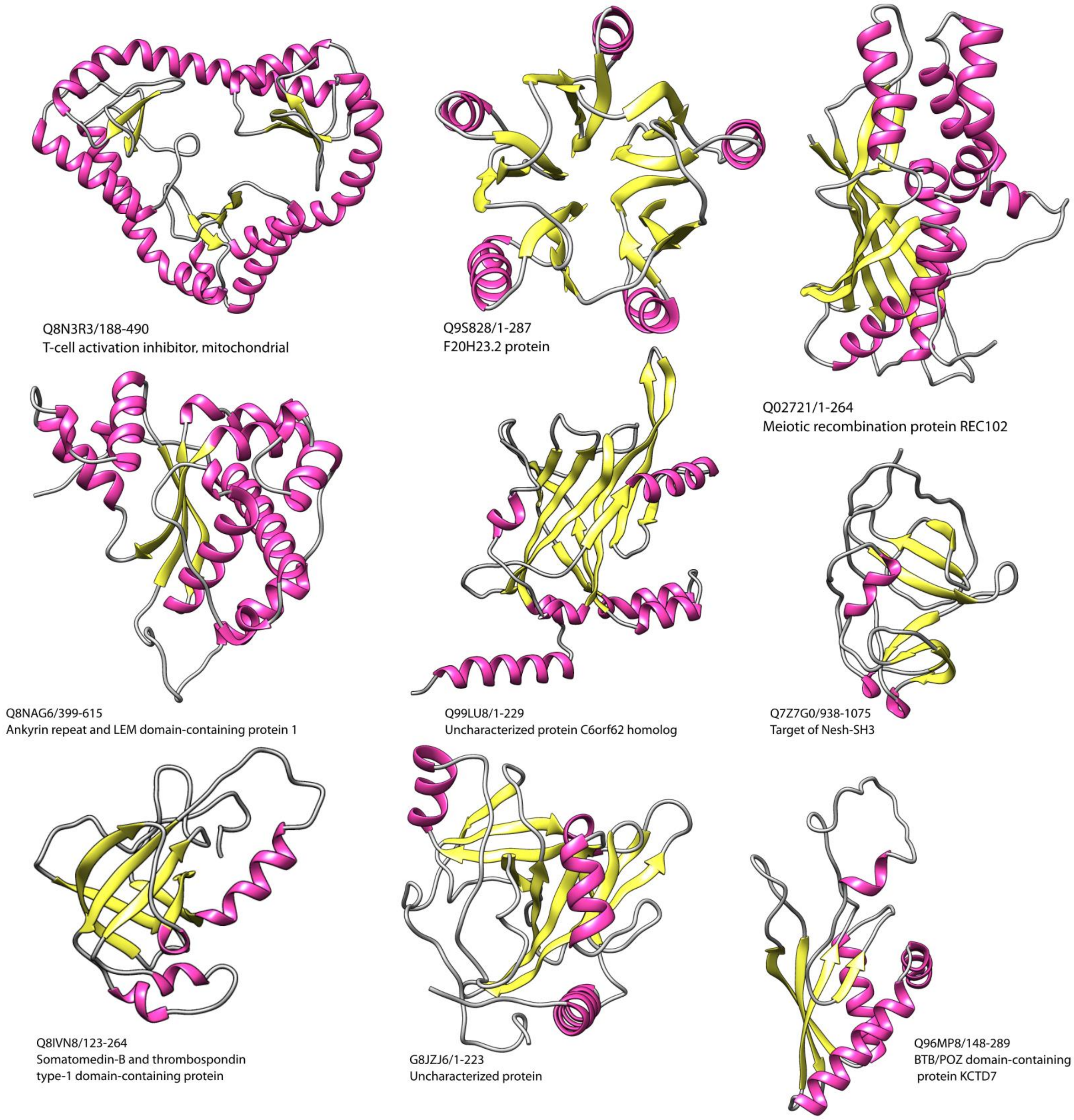


**Supplementary Figure 7.** All alpha/beta novel folds in AF2.

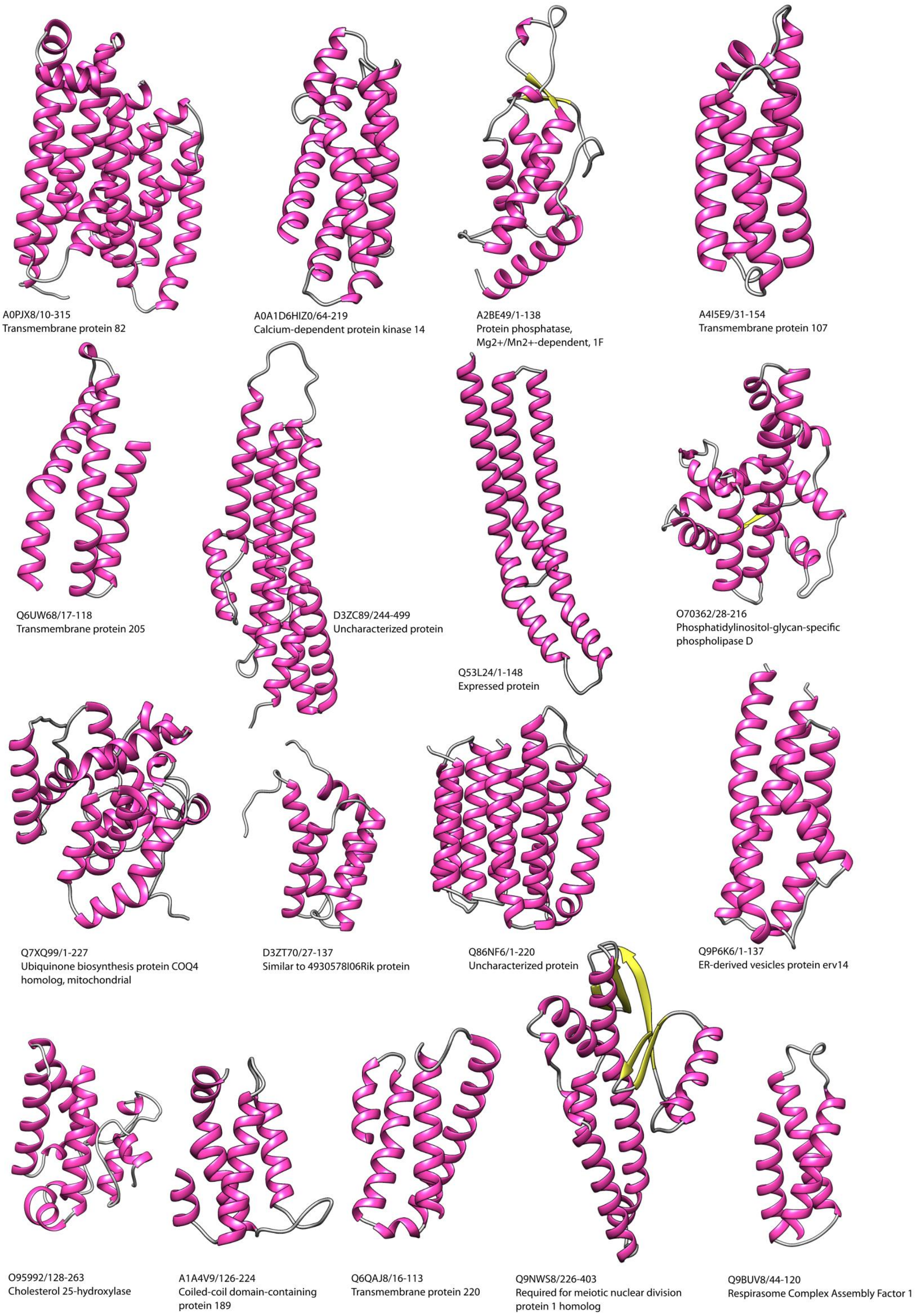


**Supplementary Figure 8.** All mainly-alpha novel folds in AF2.

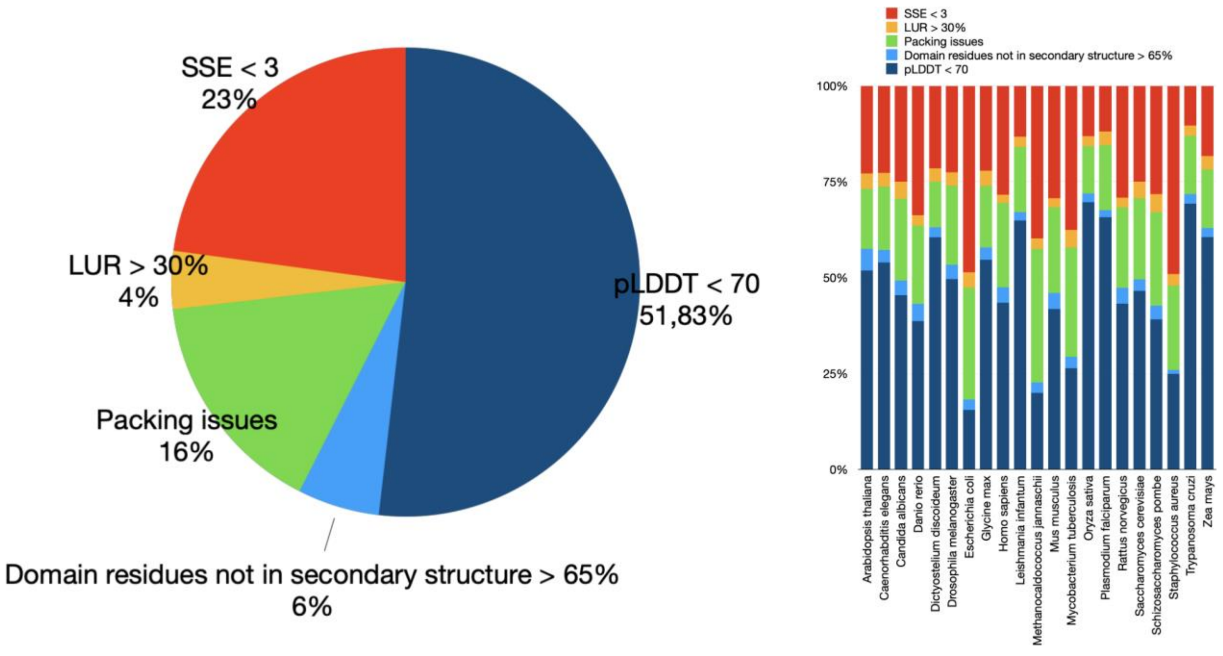


**Supplementary Figure 9.** Distribution of AF2 domains not included in CATH (left) by organism (right).

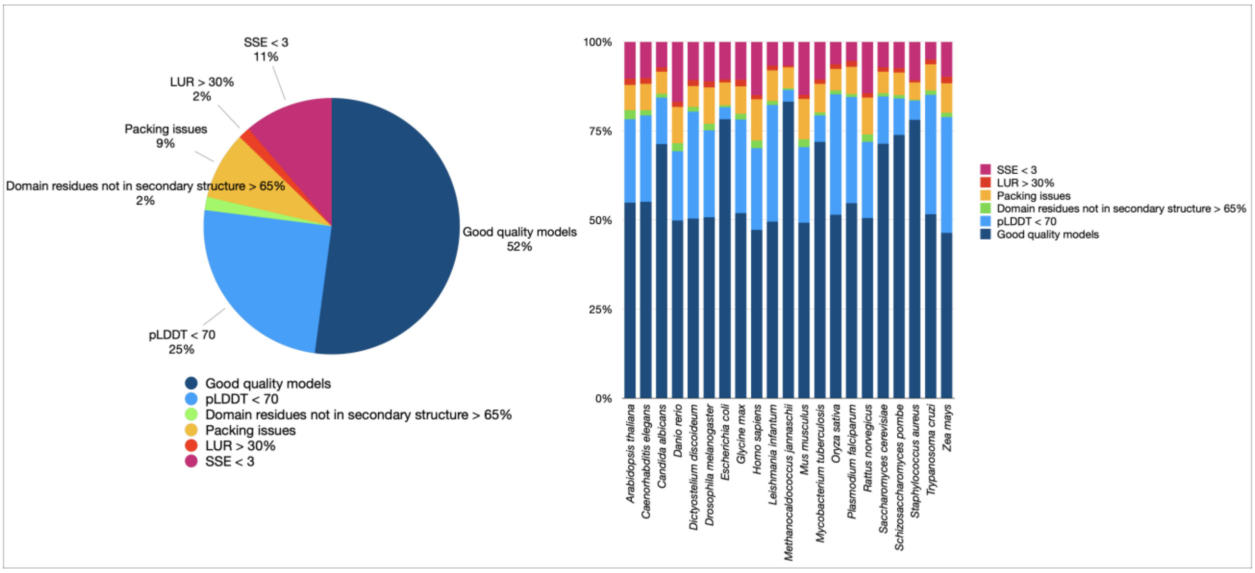


**Supplementary Figure 10.** Distribution of AF2 domains with good model quality and discarded (left) by organism (right).

**Benchmarking SSAP and Foldseek for homologs detection**

To assess the score thresholds for homologs detection using SSAP and Foldseek, we created a dataset of 3,186 curated domains that are S30 representatives of CATH that are equivalent in the SCOP classification. As the relationship of each pair of domains is known, we created an all-vs-all half-matrix of structural comparisons to be run using Foldseek and SSAP. The half matrix of pairwise comparisons consists of 13,443 homologous pairs, 67,917 pairs that share the same fold and 4,992,345 non-homologous pairs.

**Foldseek Benchmark**

We ran Foldseek (version 6315e9b67d08fb7867d6573d38d473a5b01e365d, 06/01/22) using a sensitivity threshold equal to 9 (highest sensitivity, personal communication from Foldseek developers) and retaining as many hits as possible in order to create the half-matrix. Results were parsed and an overlap based over the length of the longest structure was calculated. If a pair was missing from the final output, we included it in the results with bitscore and overlap set to zero.

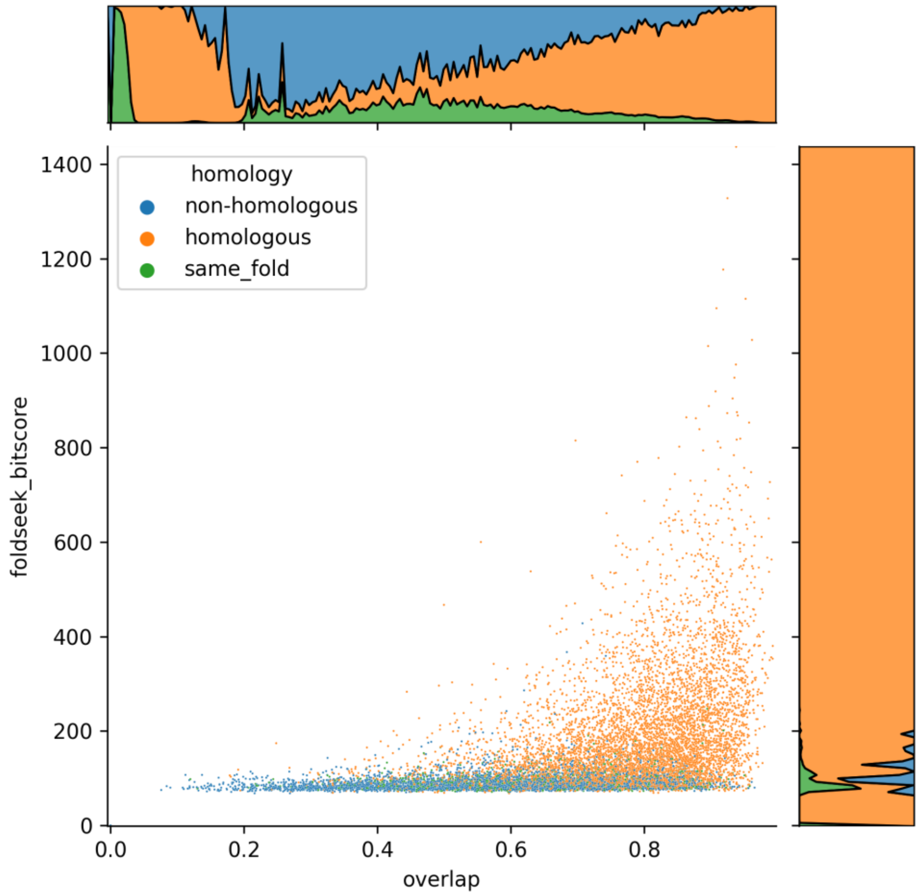


**Supplementary Figure 11:** Foldseek bitscore plotted against the structural alignment overlap. Each pair of comparisons was coloured according to their homology.

In order to calculate a homology threshold for each CATH class, we divided the dataset into pairs where both query and target belonged to the same CATH class, and calculated the percentage of non-homologous pairs over the total number of non-homologous pairs at a threshold of 60% overlap for all bitscores in the dataset. We identified bitscore cut-offs for homology at 5% error rate at 116, 165 and 117 for Class 1, 2 and 3 respectively.

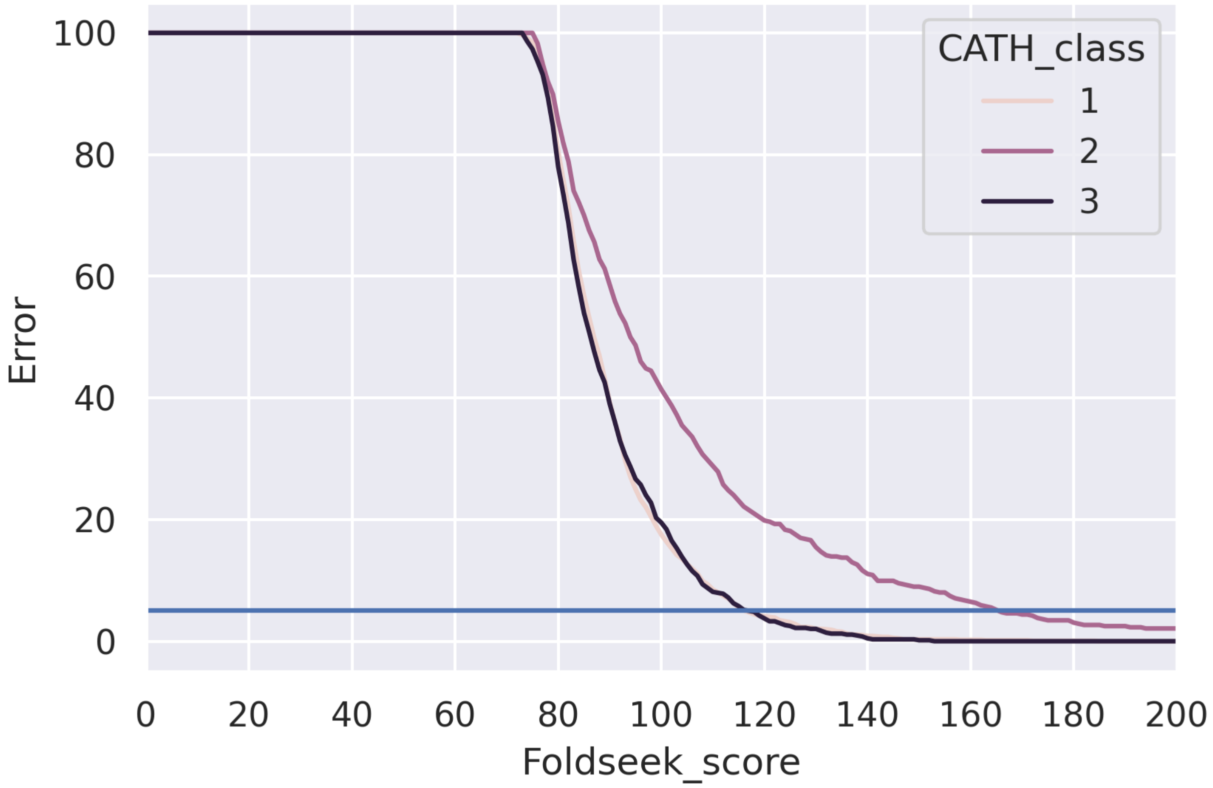


**Supplementary Figure 12:** Error rate by Foldseek bitscore for each CATH class. The horizontal blue line represents the 5% error threshold.

**SSAP Benchmark**

All domains in the S30 dataset were scanned in an all-vs-all fashion using SSAP. Since SSAP performs pairwise comparisons, one for each run, the half-matrix was generated directly without requiring additional missing pairs in the output.

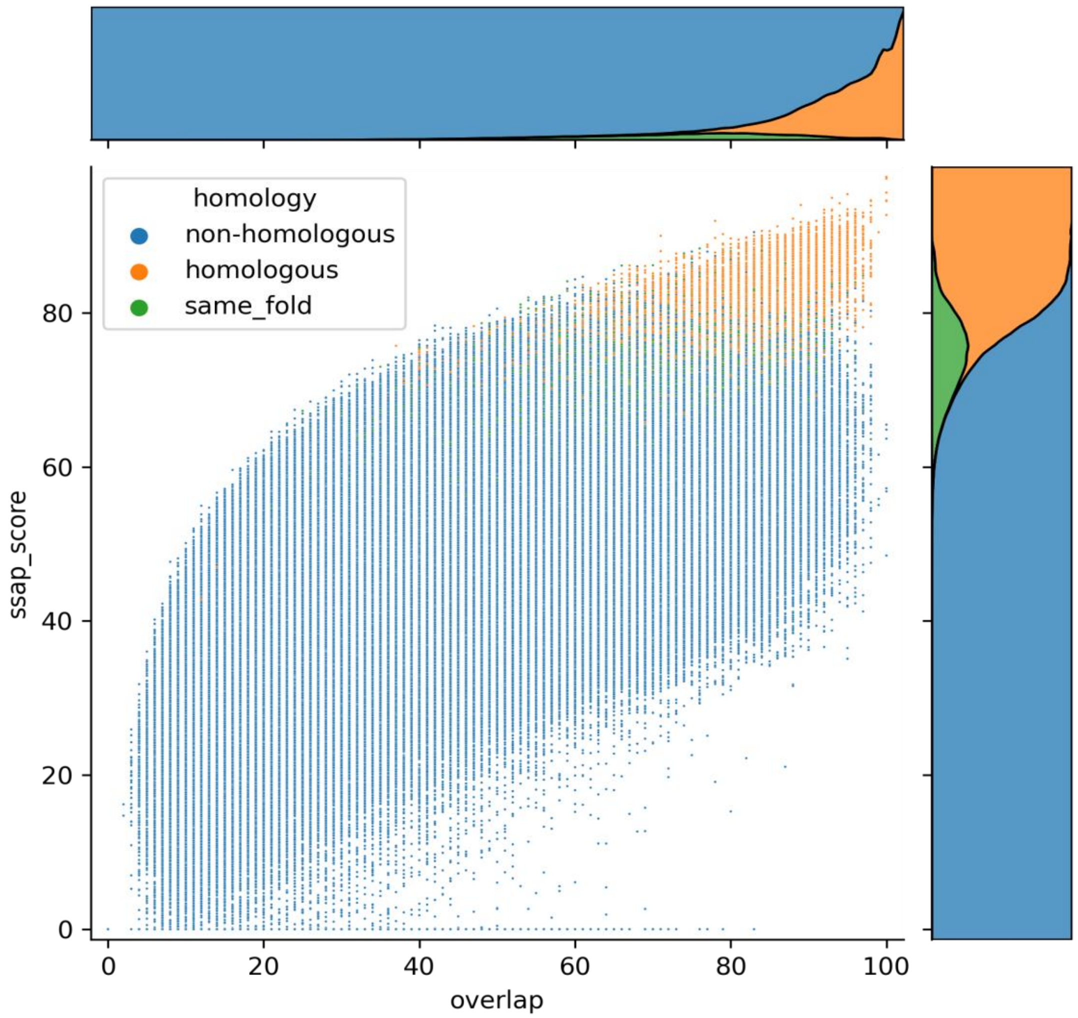


**Supplementary Figure 13:** SSAP score plotted against the structural alignment overlap calculated as 100% x overlap /length of largest domain.

Each pair of comparisons was coloured according to their homology.

The error rate was calculated in the same fashion as the Foldseek benchmark, resulting in a SSAP score threshold at an overlap of 60% of 71, 66 and 69 for CATH Class 1, 2 and 3 respectively.

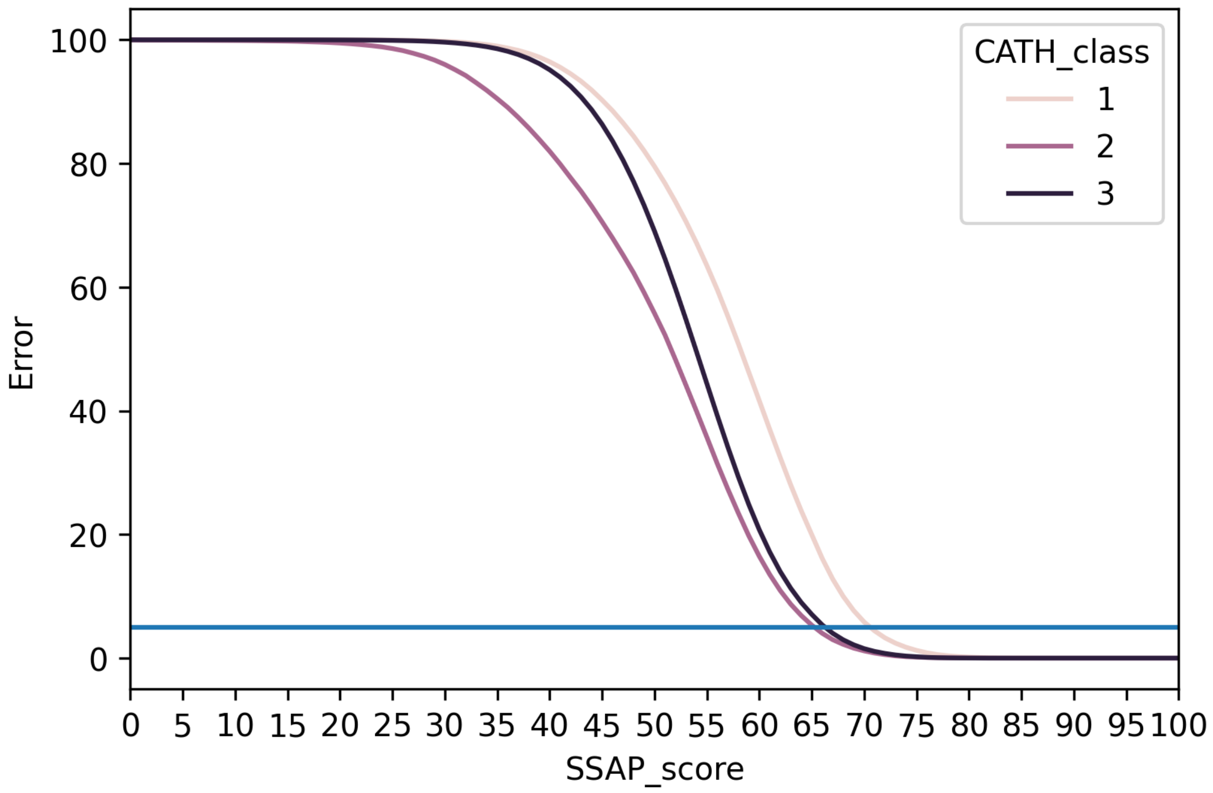


**Supplementary Figure 14:** Error rate by SSAP score for each CATH class. The horizontal blue line represents the 5% error threshold.
